# Synapse-based bispecific immune cell engager model predicts invariance in synapse behavior across different effector-to-tumor cell ratios

**DOI:** 10.64898/2026.06.24.733437

**Authors:** Michael W. Chevalier, Zhiwei Zhang, John Tolsma, Michael G. Zager

## Abstract

Immune cell engagers (ICE) such as bispecific antibodies (bsAbs), within an immunological synapse, bind and link CD3 on a T cell to a target antigen (TAA) on a cancer cell, forming a trimer (CD3:bsAb:TAA complex). With sufficient trimer numbers within the synapse, the T cell can become activated and promote cancer cell killing. Elranatamab, a CD3-bispecific antibody for multiple myeloma, has received FDA and EMA filing acceptance (August 2023 and December 2023, respectively) adding to a growing list of bsAbs that are treating patients. In the drug development stages of ICE bsAbs, mechanistic modeling approaches are often used to attain a greater quantitative understanding of the modality, preclinically, and provide human pharmacokinetic and efficacious dose predictions to aide in Phase 1 trial design. To date, the majority of ordinary differential equation (ODE) trimer models treat the tumor compartment as well-mixed and trimer formation is governed by a bulk population reaction not accounting for individual synapses. This lack of discrimination can lead to imprecise analysis when analyzing results across E:T ratios using metrics like trimers per T cell or trimers per target cell. To this end we developed an ODE trimer model based on single-synapse complexes (one target cell/one immune cell) with 2D cross-linking trimer formation. We show computationally that the number of trimers per synapse is invariant to the value of the E:T ratio for a given free bsAb concentration, a property that cannot be captured by non-synapse models. A simple demonstration of this discrepancy using the well-known Betts trimer model is presented. We then apply the Betts trimer model coupled to a tumor growth inhibition (TGI) module to show that our synapse-based trimer model is easy to substitute in to model TGI, including the addition of a trimer-per-synapse activation threshold function for cell killing. Overall, our model attempts to balance mechanistic fidelity while limiting the complexity of the model.

## Introduction

Immune cell engagers (ICE) such as T-cell bispecifics [1, 2, 3] and other immune-cell-type bispecifics [4] are at the forefront of immuno-oncology. ICE has found success for liquid tumors, such as multiple myeloma, which includes BCMA-targeted bispecific therapies, including teclistamab and elranatamab [5]. Solid tumors have been shown to be more of a challenge for ICE, due to the immunosuppressive Tumor Microenvironment (TME) [4, 6, 7, 8] where ‘exhaustion’ of activated T cells can occur as well as insufficient trafficking of T cells into the solid tumor to enable tumor cell killing. Making therapeutic progress to overcome these barriers is critical and an area of intense research. In the preclinical space, there have been promising studies of solid tumors which have provided proof of concept for bispecifics such as targeting P-cadherin [9], prevalent in pancreatic, gastric, and colorectal cancers. For many of these programs, Quantitative Systems Pharmacology modeling has been invaluable, providing insight into how the therapy affects tumor growth, in vitro-to-in vivo translation, and human predictions [10, 11].

The modeling approaches of ICE bispecifics have mostly focused on well-mixed trimer models [10] to capture the bispecific interactions between T-cells and target cells in the tumor by forming trimers in a well-mixed compartment to bridge the two cell types, analogous to the T-cell TCR binding to an antigen on a target cell [12]. However, one mathematical problem is that individual cells and synapse pairs, for example, are not accounted for. This effectively allows all immune cell antigens (IAAs) to react with all target cell antigens (TAAs) in the space (see diagram in Figure 1A). These reactions are effectively treated as bulk 3D reactions [10] even for the trimer formation, with some approaches applying an avidity reaction *χ* factor [13, 14] for the final step of trimer formation [15, 16] but still assuming a 3D reaction volume. Most recently, this work [17] was extended for having trimer formation occur as a 2D avidity reaction to account for the 2D diffusion cross-linking reaction occurring on the membranes. The researchers transform the reaction space volume to a 2D space accounting for all immune and target cells. This approach, while keeping the model relatively simple, also demonstrates data fitting capabilities that exceed prior models. However, individual synapses are not modeled, only trimer formation is. This can be a potential limitation when trying to relate trimer concentrations to the trimer numbers at the single immune cell level using metrics like ‘trimers per effector cell’. For synapse models, one uses ‘trimers per synapse’, a more biologically relevant metric given that it is formation of the synapse that allows for target cell killing.

**Figure 1:**
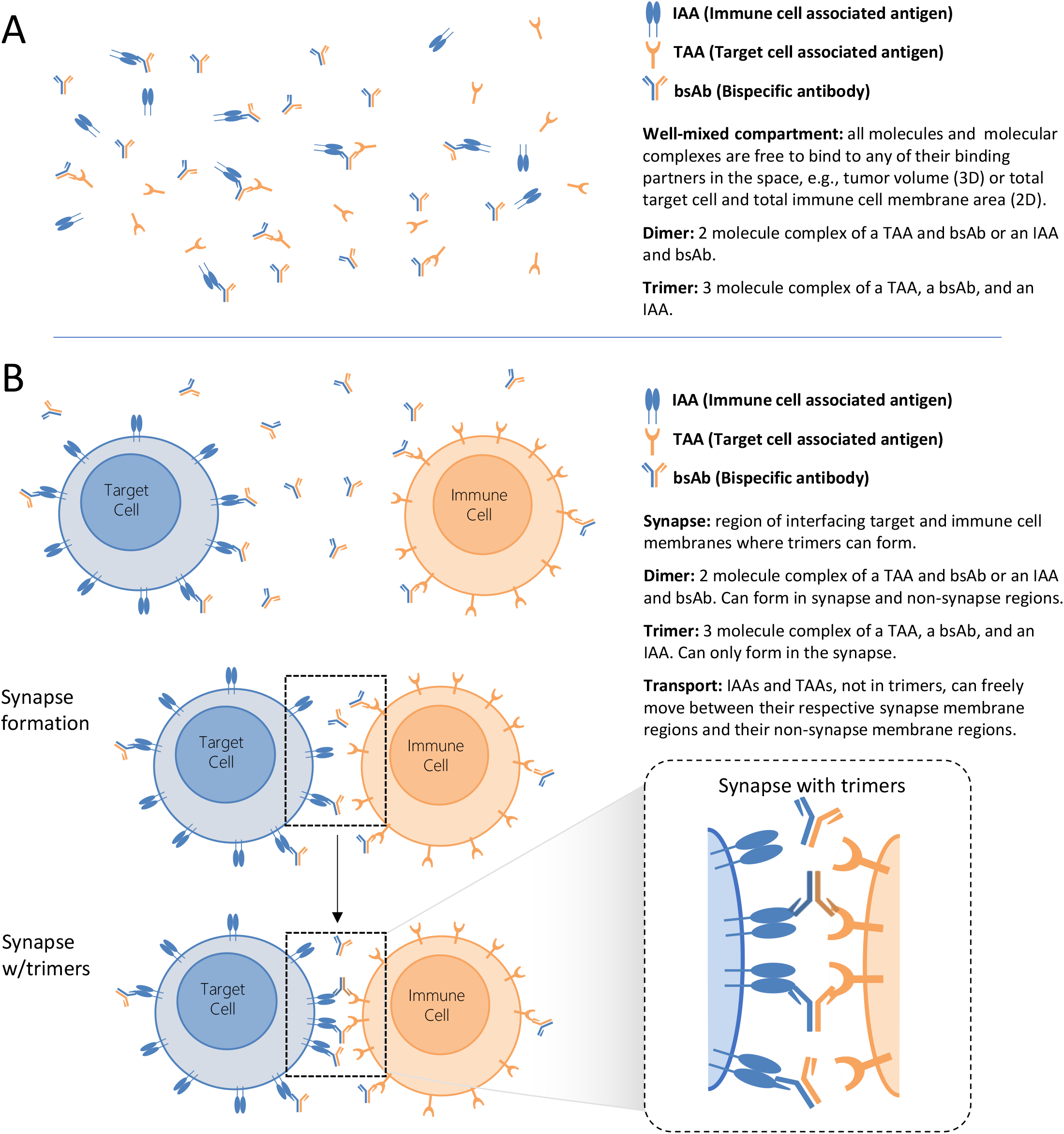
Differentiating between a well-mixed trimer model and a basic synapse model. (A) Process of trimer formation in a well-mixed compartment. (B) Process of synapse formation with bsAb induced trimer connections that bridge together an immune cell and target cell. Synapse model illustration shows an immune cell and target cell (top two cells) that come together to form a synapse (middle two cells), i.e., an interface of their respective cell membranes. Within the synapse, when the bispecific antibody is present, trimers can form between the immune cell and target cell (bottom two cells). With sufficient numbers of trimers, this can lead to immune cell killing of target cells.

One observation of these well-mixed methods is that as the effector cell to target cell ratio (E:T ratio) changes and the bsAb concentration remains the same, the number of trimers per effector cell changes. While this observation is not necessarily striking, the limitations of a model that does not differentiate which immune cells are engaged to target cells and which aren’t lends itself to interpretation that is not necessarily grounded in known biology, i.e., the existence of sub-populations of synapse complexes, free T cells and free target cells. Now if synapses are modeled, there is a prediction that the number of trimers per synapse should not change across different E:T ratios for a given free bsAb concentration.

Recent work does model the synapse applying agent-based modeling [18], where individual synapses are formed and modeled over time and where trimers can form based on a diffusion-reaction term that cross-links the bispecific to the IAA and the TAA. Their approach follows individual cells forming synapses. For our work, we developed an ordinary-differential-equation (ODE) trimer model based on synapses, focused on modeling single-synapse complexes (synapse complex comprised of one immune cell and one target cell) as an initial step towards modeling more general multi-synapse complexes (synapse complex comprised of at least 3 cells with at least 1 immune cell and 1 target cell, yielding 2 or more synapses [18]). Our work explicitly models synapses where trimer formation is based on an avidity cross-link reaction within the synapse itself. To achieve this we developed a simple, seamless approach that partitions the bulk synapse concentration for each reacting species in the avidity reaction to the single synapse level. The avidity reaction term then scales back up to solve for the total concentration of trimers over time. This innovation is effectively a partitioning approach to ensure the proper number of molecules are available for reacting at each synapse. Prior ODE approaches do not account for this, which we will show is crucial for consistent calculations of trimers across changing numbers of effector cells. In vivo, effector cell numbers over time can span many orders of magnitude. Recently, an ODE synapse model [19] has been developed. However, rather than bulk populations which we focus on, their approach applies ODE-dependent trimer formation for individual synapses that form from the populations over time, somewhat similar to the agent-based synapse modeling approach describe above [18] developed by the same research group.

A major motivation for our synapse approach is not just to have a more structured and consistent trimer calculation, but it also allows one to model which immune cells are engaged in synapses and which ones aren’t. From this structure we can naturally add different immune cell types that would be in the tumor and model the interactions between these various cell types, if we so choose or if data suggests the need. In other words, we can develop a fit-for-purpose model that reflects the data with the synapse model helping to serve as the foundation to build upon. Furthermore, because we are modeling the number of trimers formed at each synapse, this allows us to set thresholds of immune cell activation for synapses, e.g. minimum number of trimers required for activation. Overall, with slightly more model complexity, our approach has the potential to yield a more realistic model with more flexibility.

We present a simple demonstration of discrepancies between the synapse model and older models across E:T ratios. Using the well-known Betts trimer model [10], we calculate the trimers per T cell whose magnitude varies over E:T ratios, while for the synapse model, the trimers per synapse magnitude is invariant. We then simulate the Betts trimer model coupled to a tumor growth inhibition (TGI) module to show that our synapse-based trimer model is easy to substitute in and simulate TGI with it.

Finally, because our synapse model is general, we can easily adapt the approach to other immune cell therapies such as Chimeric Antigen Receptor T cells (CAR T) as well as general immune cell interactions that form synapses with target cells. To this end, we present a demonstration of CAR-T modeling as well as applying an antibody to block an immune cell/target cell interaction such as anti-PD1 or anti-PDL1 therapy [20].

Overall, our ODE synapse model attempts to balance mechanistic fidelity while limiting the complexity of the model.

## Results

### Trimer formation in the synapse: how membrane-bound bimolecular reaction propensities scale from the single synapse case to the overall synapse population

When an immune cell and target cell come in close proximity, they can form a region known as an immune synapse (diagram in Figure 1B). In the synapse where their cell membranes and membrane-bound receptors are in close proximity, a bispecific antibody (bsAb) can form a trimer between a target associated antigen (TAA) on the target cell and an immune cell associated antigen (IAA) on the immune cell. Only in the synapse can this occur. The trimer is formed in two possible ways. In the synapse, a dimer of a TAA and a bsAb binds to an IAA or a dimer of an IAA and a bsAb binds to a TAA. Dimers can form in both the synapse and non-synapse regions. IAAs and TAAs that are not bound as trimers can freely move (via diffusion-driven transport) between their respective synapse membrane regions and their non-synapse membrane regions. Generally the number of trimers in the synapse drive the level of immune cell activation (corresponding to the level of target cell killing) which can be a nonlinear function of the synapse trimer number [11]. Integral to this, a dependence on the geometry of the bsAb binding such as membrane-proximal epitopes have been observed to enhance cell killing [21]. Thus, epitope choices can be critical to achieve good bsAb designs.

We will now go through the mathematics of the reaction steps to form trimers. Along the way, we will highlight and discuss important differences in mathematical structure between the two bimolecular reactions that occur in this stepwise process of forming trimers. For each dimer formation reaction on single cells (target cell or immune cell, alone or in synapse complexes), we will start with the single cell reaction rate and scale by the population number per tumor interstitial volume which will yield the familiar bulk bimolecular reaction rate in the tumor. However, when using the same approach for trimer formation in the synapse, which involves both the target cell and the immune cell, this will yield a different mathematical formulation.

The first reaction that can occur in the formation of the trimer is for the free bispecific antibody in the tumor (*bsAb*_*t*_) to bind to either the immune cell associated antigen (*IAA*_*t*_) or to the target cell associated antigen (*TAA*_*t*_) in the free target cells and free immune cells (top diagram of Figure 1B), as well as the target cells and immune cells in synapse pairs (middle and bottom diagrams of Figure 1B). We will occasionally refer to these various dimers as bsAb-dimers. Crucially, we’ll start at the single cell level, with the reaction rate contribution from a single cell, and scale to the population concentration level within the tumor. For free target cells, let {*TAA*_*tsc*_*}* be the number of free TAA per target cell in nmol per cell (sc stands for single cell) at any given time. Here the expression for the overall concentration of *TAA*_*t*_ would be 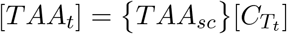 where 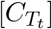 is the target cell number per tumor interstitial volume. The term for the reaction rate of TAA binding to the free bispecific antibody, given the rate constant 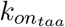 and the tumor interstitial volume *V*_*t*_, can be written as

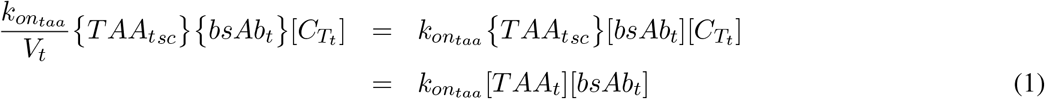

where the term on the left-hand side is just the single-cell reaction rate (nmol per time, per cell) scaled by the number of cells per volume 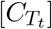 to yield the total reaction rate (nM per time). On the left side, {*bsAb*_*t*_} is the free bispecific antibody in the tumor in nmol representation. This left side of the equation equates to the familiar 3D bimolecular reaction term on the right-hand side. For target cells in synapse complexes, let {*TAA*_*sc*_*}*_*syn*_ and {*TAA*_*sc*_*}*_*non*_ be the number of free TAA per target cell in nmol per synapse complex at any given time in the synapse region and non-synapse region, respectively. We define 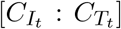 as the number of synapse complexes per tumor interstitial volume. Note that this also equal to the number of synapses per tumor interstitial volume since we are modeling single-synapse complexes in this work. Following the logic of Eq. (1) starting from the left-side, but scaling by 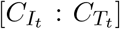 instead, their respective reaction rates would be 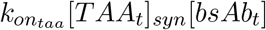 and 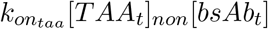.

Following the same approach for the immune cell associated antigen on free immune cells, the reaction rate term for IAA binding to a free bispecific antibody would be

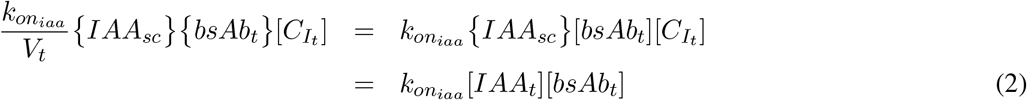

where 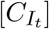 is the immune cell number per tumor interstitial volume. Similarly for the immune cells in synapse complexes, let {*IAA*_*sc*_*}*_*syn*_ and {*IAA*_*sc*_*}*_*non*_ be the number of free IAA per immune cell in nmol per synapse complex at any given time in the synapse region and non-synapse region, respectively. Following the logic of Eq. (2) starting from the left-side, but scaling by 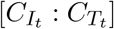 instead, their respective reaction rates would be 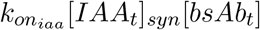 and 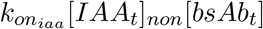.

If we set aside trimer formation for a moment, the bsAb-dimer reactions above produce the same number of total bsAb dimers in the target and immune cell populations, free and in synapses, as seen in previous trimer models for any given bsAb tumor concentration [10, 15, 16, 17]. That is, the total bsAb-dimers across all target cells and all immune cells over time will be the same across these various models. Because of this, we can strictly differentiate our model from others in how the trimers are mathematically formed, i.e. through inclusion of a synapse model. What we are doing different is two fold. One is that for immune cell/target cell synapse pairs, we define a synapse region and non synapse region for both the immune cell and target cell. This allows transport of IAAs and TAAs, not in trimers, between the synapse and non-synapse regions as discussed in Figure 1B. The other distinction is that we only allow trimers to form from molecules that are in the synapse, i.e. local. We will first discuss the mathematical form of trimer formation and later we’ll mathematically define our transport model.

For each of the bsAb-dimer reaction rate terms defined above, one of the molecules is diffusing within the solution, i.e., free bsAb, while the other one is on the surface of their associated cell type. In contrast, for the formation of the trimer in synapses (middle and lower illustrations in Figure 1B), both molecules involved in the reaction to form the trimer are on their associated cell membranes within the synapse. To restate, to form the trimer in the final step, the reacting molecules are either an IAA-bsAb dimer and a TAA or a bsAb-TAA dimer and an IAA.

In either case, the reaction step to forming a trimer involves 2D diffusion of the reacting molecules on their respective membranes until a cross-linked binding event occurs. The rate constant for this process we denote as 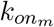 where ‘m’ stands for membrane. *A*_*syn*_ represents the membrane area of the synapse (the same for both cells at the synapse). Thus, the term for the reaction rate of a TAA binding to a bispecific antibody bound to an IAA (IAA-bsAb dimer) can be written as

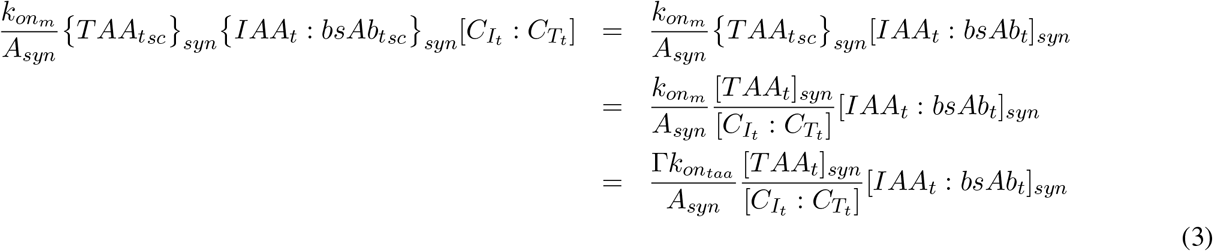

where 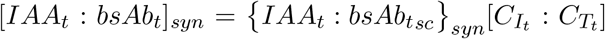 and ‘sc’ represents single cell. We also use the relationships 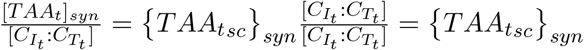. The left-hand side of Eq. (3) is the single synapse reaction rate (nmol per time, per synapse complex), taken from [17] (Eq. S1.1 in Supplemental Materials), and scaled by the number of synapse complexes per tumor interstitial volume, 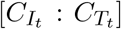. Dimensionally, the left-side yields nM/time. We represent 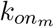 by 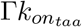 where 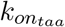 is the typical IgG1 3D association rate. While there are theoretical formulas for 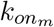, they require biophysical measurements of various diffusion rates as well as how accessible the arms of the particular bsAb are to bind and form a trimer [22, 23]. Thus, due to these difficulties, we will typically fit Γ. For this paper, the membrane area of the synapse on both the target and the immune cell we will assume to be *A*_*syn*_ = 7.854 *×* 10^−11^ m^2^ or 78.54 *µ*m^2^ [24]. This assumes a circular synapse cross-section with a radius of 5 *µ*m, within the typical range for an immunological synapse [24].

Similarly the term for the reaction rate of IAA binding to a bispecific antibody bound to a TAA (bsAb-TAA dimer) can be written as

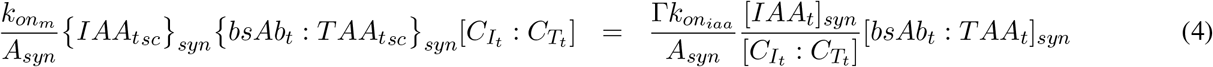

Our derivation of the trimer-formation reaction rates in Eqs (3) and (4) result in 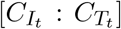 in the denominator, which is crucial when the number of synapses changes over time. To date, the trimer formation terms in various models within the field do not contain this. We will later show that this produces errors when this synapse density term is missing.

In summary, for the form of the trimer bimolecular reaction rate above (final right hand side of Eqs (3) and (4)), the scaling from single cell (nmol) to total synapse concentration results in a synapses-per-volume term, i.e., 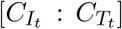, in the denominator. This term ensures the correct number of reacting molecules per synapse. While for the bimolecular reactions that involve one species on a cell and one in the medium, e.g. a free mAb, the scaling from the single cell (nmol) to total population concentration results in the typical 3D bimolecular reaction rate that we are accustom to (Eqs. (1) and (2)).

### Transport into and out of the synapse membrane region for cells in synapse complexes

We will first consider immune cell/target cell pairs that form a synapse. Here we derive their transport rate relationships of receptors, with or without bsAb bound, between the synapse membrane region and the non-synapse membrane region for both target and and immune cells. We will derive it for receptors only, but the same form of the derivation can be applied to receptors bound to bsAb. For each cell type, we assume a compartment for the synapse and a compartment for the non-synapse region with exchange rates between the compartments. In addition, we also enforce simple physical constraints based on the 2D membrane environment. As discussed above, at a given synapse, the surface areas of the target cell membrane interfacing with the immune cell membrane are assumed to be the same, i.e., *A*_*syn*_. For the target cell’s membrane in the non-synapse region, we denote its surface area as 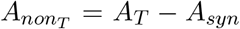 where *A*_*T*_ is the total area of the target cell membrane. Likewise for the immune cells, 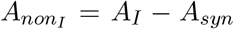 where *A*_*I*_ is the total area of the immune cell membrane.

Physically speaking, for the target cell, we assume the TAAs diffuse on the membrane in and out of the the synapse. For this simple diffusion model, when there is no bispecific antibody present, at steady state the 2D concentration (TAA number divided by surface area) must be the same in the synapse and non-synapse membrane regions. That is,

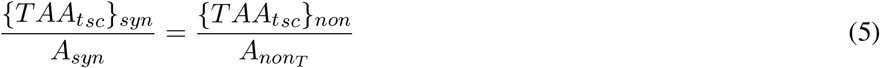

where as above {*TAA*_*tsc*_*}*_*syn*_ is single cell TAA number in nmol in the synapse. {*TAA*_*tsc*_*}*_*non*_ is the same but for the non-synapse region. Borrowing from the derivation of Eq. (3), Eq. (5) can be written as

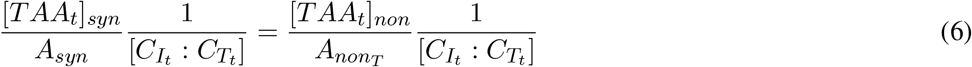

where we represent the surface concentration with the 3D volume concentrations [*TAA*_*t*_]_*syn*_ and [*TAA*_*t*_]_*non*_ divided by the number of synapses per volume. Now for our simple compartment model approach, the time-dependent transport equation for [*TAA*]_*syn*_ when no bsAb is present would be

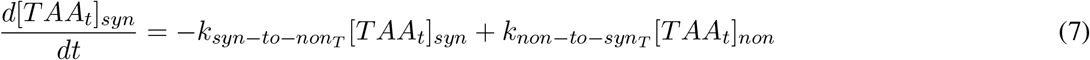

where 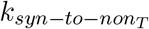 is the TAA transport rate from the synapse TAA compartment (signifying synapse membrane region) to the non-synapse TAA compartment (signifying non-synapse membrane region), and vice versa for 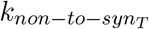. Note that the time-dependent transport equation for [*TAA*_*t*_]_*non*_ would have the same right-hand side but with opposite signs. At steady state Eq. (7) becomes

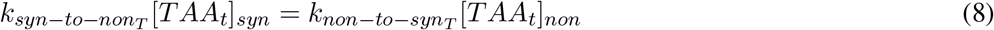

Well now derive constraints on the relationship between 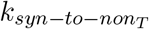 and 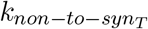. Now in order to satisfy the physical constraint that the 2D membrane concentrations be equal in the synapse and non-synapse membrane regions, this requires that

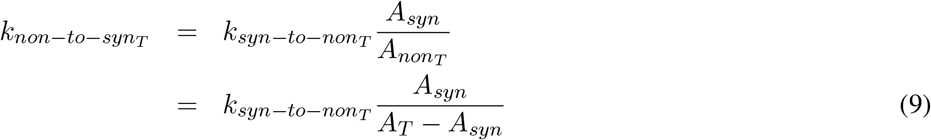

where we get the expression in Eq. 9 by dividing Eq. 8 by Eq. 6 and solving for 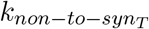. We’ll also assume that 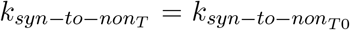 the transfer rate from the synapse to outside the synapse when only a single synapse is on the target cell. The number of synapses can be larger [18], but for this paper we only consider a single synapse on the target cell. Following the above logic, the immune cell involved in the synapse would have the transport rate relationship

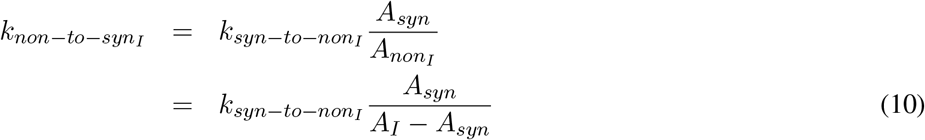

where we’ll also assume that 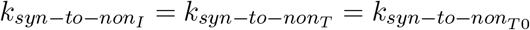.

The timescales of transport should coincide with that of transport by diffusion across the surface area of the cell membrane of the cell. We’ll assume an upper diffusion coefficient of membrane receptors to be 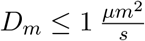 [25, 26]. For T cells which have a maximum surface area of 200 *µm*^2^, the time to spread from the synapse into the non-synapse region of the membrane would take around 20 to 200 seconds, approximately. For the purpose of this paper we’ll test rates that correspond spread times of about 6s to over 2 hours and show that our results are invariant over this range. Essentially, as long as the ratio relationships at steady state hold, i.e. Eq. 9, the specific values of 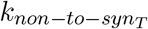 and 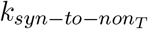 essentially do not matter as far as the levels of steady-state trimer formation is concerned.

As mentioned above, for receptors bound to bsAb (immune cell or target cell) they will have the same relationships derived above for the receptor only cases, but the magnitudes could be different, maybe slightly slower because the bsAb is attached. However, for the simulations in this paper, we assume they have the same magnitude.

### For a given bsAb tumor concentration, the single-synapse model predicts the number of trimers per synapse is invariant across different E:T ratios and a large range of biologically relevant transport timescales

A repercussion of the form of the trimer bimolecular reaction in Eqs (3) and (4) is that at a given steady state tumor bsAb concentration, each synapse is independent of the other synapses. Therefore, the number of trimers per synapse will be independent of the number of synapses. Another way to think about this is it will be independent of the effector-to-target ratio (E:T ratio). This is because Eqs (3) and (4) ensures that each synapse is partitioned with the proper number of reacting molecules to form trimers, regardless of the number of synapses per volume 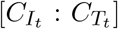. Note that in this paper, when we discuss bsAb concentration (or drug concentration) in the manuscript text and plots, we are always referring to the free bsAb concentration (or free drug concentration).

To demonstrate that the trimers per synapse will be independent of E:T ratio, at a given bsAb concentration at or near steady state, we first ran PKPD simulations with a tumor volume for different doses (for model details see Supplementary Information S3). For these simulations we set IAA levels to 100,000 receptors per cell and TAA levels to 10,000 receptors per cell, *KD*_*taa*_ = 1 nM, and *KD*_*iaa*_ = 10 nM, and Γ = 200. Note that for this paper, the IAA levels will always be 100,000 receptors per cell while TAA receptors per cell will be varied. Each simulation run had different transport rate values and E:T ratios. Figure 2 presents the dose-dependent trimer dynamics in the left plot for each E:T ratio and synapse transport rate combination (see Figure S1 for the the dose-dependent PK (plasma and tumor) for these various cases). Across the different transport rates and E:T ratios, the dose-dependent trimer dynamics behaves essentially the same (after the initial transient). That is, after the initial transient, the trimer kinetics are in quasi-steady-state with respect to the slowly varying tumor concentrations driven by the plasma PK. To demonstrate this we ran simulations where we set the tumor to a constant bsAb concentration to calculate steady-state trimers per synapse. This was repeated for a range of concentrations (right plot for each E:T ratio and synapse transport rate combination in Figure 2, black bell curve with circles). We then plotted the average bsAb concentration in the tumor vs. the average number of trimers per synapse from the PKPD simulations which agree well with the steady-state trimers per synapse (Figure 2, vertical colored line segments in the same bell plots). Importantly these results are invariant across the different transport rates and E:T ratios as predicted by Eq (3) and (4).

**Figure 2:**
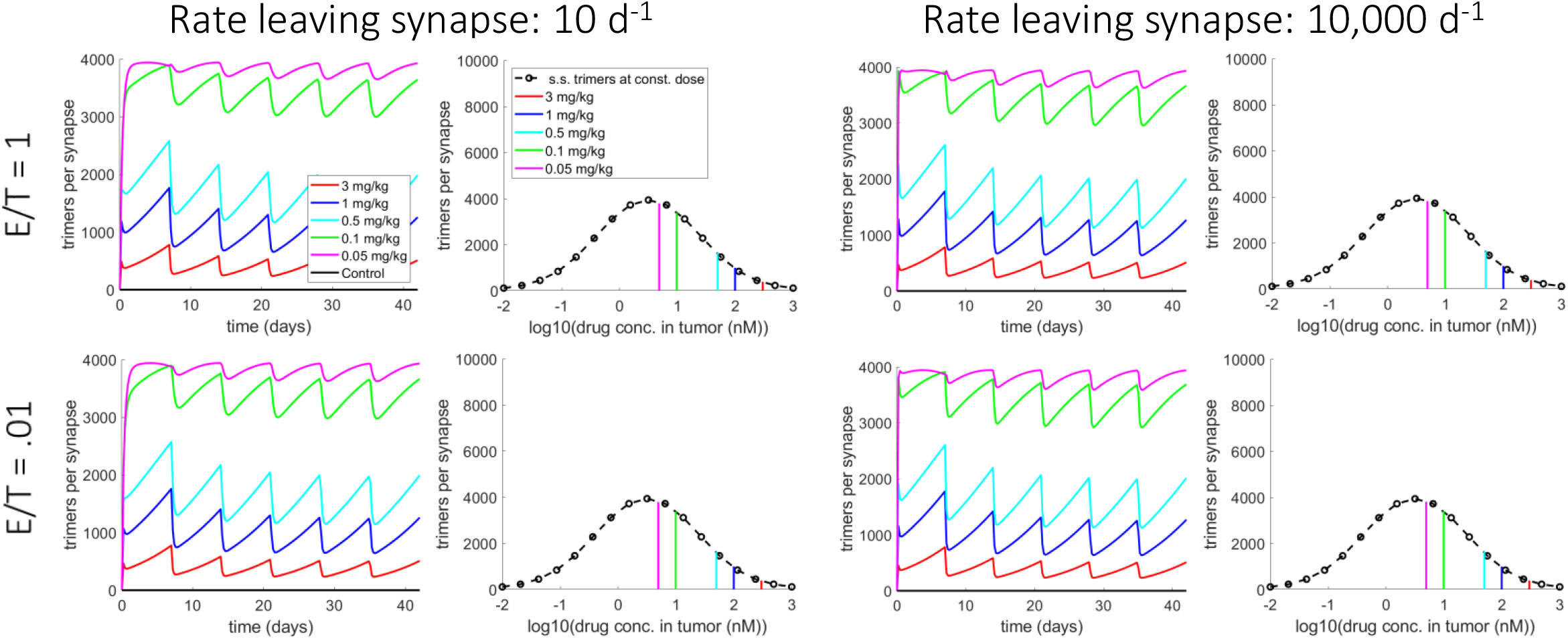
Dose-dependent trimers per synapse calculations from the synapse model exhibit invariance across different E:T ratios and transport timescales. For these simulations, the bsAb has a *KD*_*iaa*_ = 10 and *KD*_*taa*_ = 1. For the synapse transport rate leaving the synapse, simulations were run for values of 10 and 10000 d^−1^, E:T ratios of 1 and 0.01, doses of 0 mg/kg (control), 0.05 mg/kg, 0.1 mg/kg, 0.5 mg/kg, 1 mg/kg and 3 mg/kg, and Γ = 200. For each synapse transport rate and E:T ratio, the left plot presents the dose-dependent trimers per synapse over time. In the right plot, the vertical colored bars represent dose-dependent, time-averaged trimers per synapse vs. dose-dependent, time-averaged tumor bsAb concentration (see tumor PK plots in Figure S1), where ‘time-averaged’ is over the whole simulation. The black dashed-line with circles represents steady-state trimers per synapse vs. steady-state tumor bsAb concentration.

For results in the right panels of Figure 2 the plots of steady-state trimers per synapse vs. free drug concentration plots exhibit a bell-shaped curve, similar to what is observed in the Betts trimer model [10]. This is a well-described phenomenon for ternary complexes [27, 28]. When the concentrations of bsAb are low, conditions favor the formation of trimers which increase with increased bsAb concentration, initially. However, as concentrations increase even further, bsAbs will become in excess and dimers become more dominant where higher concentrations results in the further reductions in the number of trimers. Hence there will be an overall bell-shape in steady-state trimers-per-synapse vs. free drug concentration.

Next we took a closer look at the time dynamics of various molecular species on the synapse pair, not just for the trimers, but for other quantities in both the synapse and non-synapse regions of the immune cell and target cell. For the simulations that produced the 2nd row in Figure 2 (*k*_*syn*−*to*−*non*_ = 10 *d*^−1^, E:T ratio = .01, slow transport case (left two plots) and *k*_*syn*−*to*−*non*_ = 10, 000 *d*^−1^, E:T ratio = .01, fast transport case (right two plots)), we plot various time dynamic quantities driven by doses of 0 mg/kg, .05 mg/kg, and 1 mg/kg, respectively (Figure 3). For each dose dependent panel, we plot IAA-dependent quantities (top plot in each panel) and TAA-dependent quantities (bottom plot in each pannel). For IAA-dependent quantities these include [*IAA*]_*non*_, [*IAA* : *bsAb*]_*non*_, [*IAA*]_*syn*_, [*IAA* : *bsAb*]_*syn*_, and trimers [*IAA* : *bsAb* : *TAA*]_*syn*_. Where *non* corresponds to non-synapse region and *syn* corresponds to synapse region. For conservation we also plot the sum of these quantities (yellow dashed line) which should always equal total CD3 per cell, 100,000 molecules for this work. Similarly, for TAA-dependent quantities these include [*TAA*]_*non*_, [*TAA* : *bsAb*]_*non*_, [*TAA*]_*syn*_, [*TAA* : *bsAb*]_*syn*_, and trimers [*IAA* : *bsAb* : *TAA*]_*syn*_. For conservation we plot the sum of these quantities which should always equal total TAA per cells, 10,000 molecules for these particular examples.

**Figure 3:**
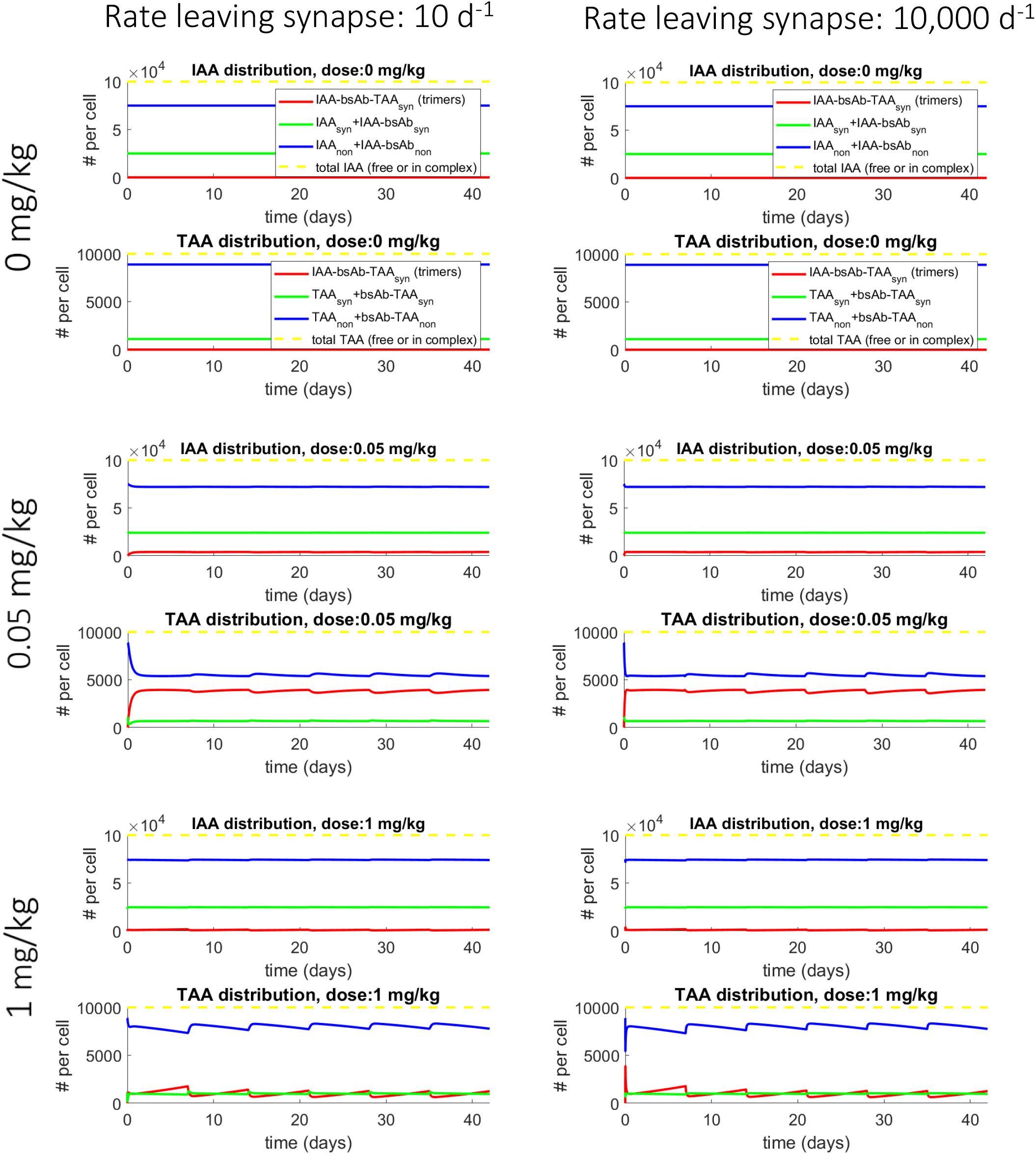
Dose-dependent dynamics of receptor transport between the synapse and non-synapse regions of the immune and target cells for different transport timescales. For the transport rate leaving the synapse, simulations were run for values of 10 and 10000 d^−1^, an E:T ratio of .01, doses of 0 mg/kg (control), .05 mg/kg, and 1 mg/kg, Γ = 200, and a *KD*_*iaa*_ = 10 and *KD*_*taa*_ = 1 for the bsAb. For each panel of a given dose and transport rate leaving the synapse, the top plot focuses on IAA receptors on the immune cell of the synapse complex. This includes plots of the total trimers in the synapse (red), the total number of IAA receptors in the synapse excluding trimers (green), and total number of IAA receptors in the non-synapse (blue), and total IAA receptors on the immune cell which is a constant (dashed yellow). The bottom plot focuses on TAA receptors on the target cell of the synapse complex. This includes plots of total trimers in the synapse (red), total number of TAA receptors in the synapse excluding trimers (green), and total number of TAA receptors in the non-synapse (blue), and total TAA receptors on the target cell which is a constant (dashed yellow).

First, at 0 mg/kg, we see no change in the quantities over time for either the slow transport case or the fast transport case. This is because the transport rates are properly matched as derived above. Otherwise, we would observe perturbations in the levels to new steady states. For the 1 mg/kg cases and the .05 mg/kg cases, the IAA plots show slight changes relative to the intial values, mainly due the fact that for this case the IAA per immune cell (100,000) is much larger than the TAA per target cell (10,000). In contrast, the TAA plots show larger relative changes in all quantities. For the 1 mg/kg cases, relatively small numbers of trimers are formed (*<* 1000, on average) with slight transport from the non-synapse region to the synapse. The fast transport case shows an initial strong drop in TAA as the concentration of bsAb in the tumor increases quickly over time and trimer levels track (approximately) along the bell-shaped curve (rising at low bsAb concentrations, dropping at higher ones) as the tumor bsAb concentration closely mirrors that of the plasma. For the slow transport case, transport is too slow for the trimers to track the bell-curve due to the tumor bsAb concentration dynamics. Interestingly, the mg/kg case is near the peak of the bell in trimer numbers (Figure 2) and the total TAA in the non-synapse region has decreased by a factor of almost two (Figure 3). Given these dose dependent changes in TAA in the non-synapse region, it is possible that we might be able fit model parameters (or determine bounds) including Γ. This could potentially be done with in-vitro imaging experiments of bsAb dose dependent synapse and non-synapse fluorescence of tagged TAA. Sampling over dose would allow us to experimentally sample the bell-shaped curve, if there is one.

Finally, to be more thorough we tested our model for an even larger range of parameters and again ran both the PKPD and steady state models for comparison. We then plotted the average bsAb concentration in the tumor vs. the average number of trimers per synapse from the PKPD simulations which agree well with the steady-state trimers per synapse vs. steady-state bsAb concentration (Figures S2, S3, S4). As a final single summary figure (Figure S5), we present the steady-state trimers per synapse vs. steady-state bsAb concentration over the entire range of parameters tested. The main point being that for a given IAA expression, TAA expression, *KD*_*taa*_, *KD*_*iaa*_, and Γ, the behavior of the steady-state trimers per synapse is invariant over the ranges tested for transport rates, and the value of the E:T ratio. This invariance as predicted is due to the form of the trimer-formation reaction rate in Eqs (3) and (4) where the synapse density term 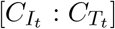 in the denominator ensures this. In the ODE formulations realm, trimer models have not had this term. Furthermore, none of these models form synapses. This is what distinguishes our formulation from other ODE formulations in the literature. However, for these other models, we can still normalize the total trimers by the number of T cells to get trimers per T cell. However, when we run these models, we will see the trimers per T cell change as we vary the E:T ratio (compare top row (model from Betts et al (2019) [10]) with middle and bottom row (synapse model) in Figure 4). This is due to the fact that individual synapses are not modeled and is a potential limitation of this approach.

**Figure 4:**
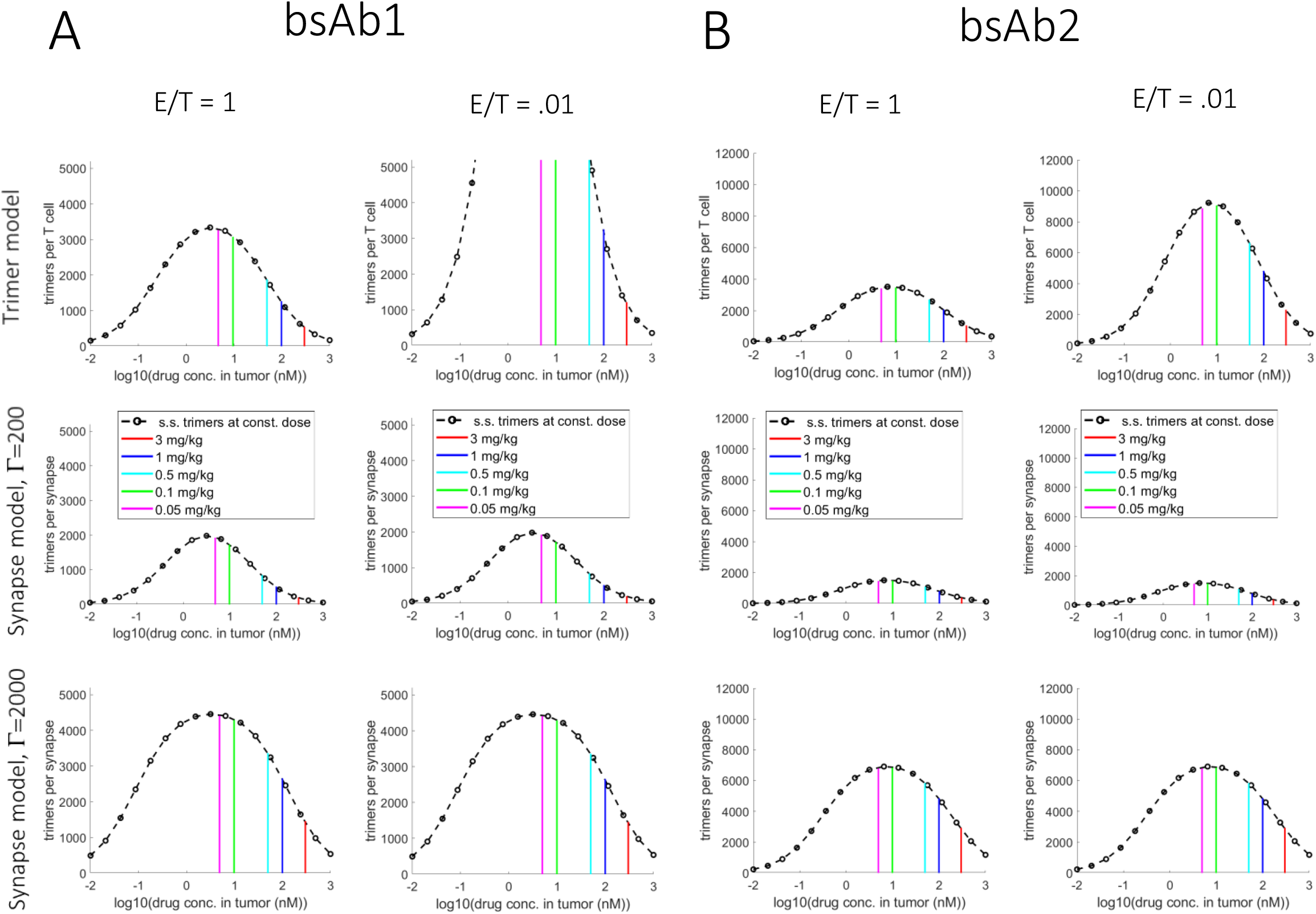
At a given average tumor drug concentration, the trimers-per-T-cell metric used for the bulk reaction trimer model [10] changes as a function of E:T ratio while the trimers-per-synapse metric used for the synapse model is invariant. A) bsAb1 (*KD*_*taa*_ = 1 nM, *KD*_*iaa*_ = 11 nM) for different E:T ratios (1 and .01) with TAA number at 5200 receptors per cell for all cases. B) bsab2 (*KD*_*taa*_ = .9 nM, *KD*_*iaa*_ = 64 nM) for different E:T ratios (1 and .01) with TAA number at 12,000 receptors per cell for all cases. For A) and B), the transport rate leaving the synapse is 10,000 d^−1^. For the synapse model, Γ = 200 (middle panels), and 2000 (bottom panels). For each plot: Vertical colored bars represent dose-dependent, time-averaged trimers per synapse (or trimers per T cell (Trimer model)) vs. dose-dependent, time-averaged tumor bsAb concentration. Black dashed-line with circles represents steady-state trimers per synapse (or trimers per T cell (Trimer model)) vs. steady-state tumor bsAb concentration.

### Inclusion of a dynamic immune cell population for transient synapse formation and other functions

The modeling above focused on trimer formation numbers for different E:T ratios. For these cases, the synapses modeled were permanent, i.e., didn’t break, which allowed us to look at steady state values. However, in reality, synapses are forming and breaking all the time, i.e., it is a transient process. The reaction rate at which the synapse forms can be modeled as a bimolecular-like reaction 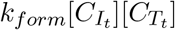 with a rate constant multiplied by the concentrations of the free immune cells and free target cells [18]. The rate of breaking (dissociation) of the synapse complex is 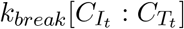. When we include this process by having a reservoir immune cell population allowed to form synapses, we will obtain a steady state population of immune cells that are in synapses, but in constant flux, because new ones are forming synapses at the same rate synapses are breaking due to the finite value of *k*_*break*_.

We will calibrate values for *k*_*form*_ and *k*_*break*_ to in-vitro synapse formation data for cell densities approaching bone marrow and solid tumors [18]. The authors observe that for effective concentrations (cells/L) of 1 *×* 10^11^ and 8 *×* 10^11^ total cells per liter (E:T = 1:1) yield about 3% and 15% of cells in synapses, respectively. To start we assumed *k*_*break*_ = 40 d^−1^ where for synapse formation rate of *k*_*form*_ = 2 *×* 10^−11^ [cells]^−1^d^−1^ this yields around 10% of cells in synapses at 4 *×* 10^11^ cells per liter at an E:T ratio of 1:1. For these values of *k*_*form*_ and *k*_*break*_, when we simulate at a lower E:T ratio such as E:T = 1:100, we find that about 20 percent of T cells are in synapses. This percentage is larger than the E:T = 1:1 case because the target cell population changes very little at an E:T = 1:100 relative to the T cell population, i.e., only 0.02 percent.

Because of the finite lifetime of a synapse, which is on the order of minutes to tens of minutes for T cell/Target cell synapses [29, 30], they will not reach as high a value of trimers per synapse as that of the permanent synapses simulated earlier (compare Figures 5A and B, full transport model) where the synapse formation rate is *k*_*form*_ = 2 *×* 10^−11^ [cells]^−1^d^−1^ and synapse break rate is *k*_*break*_ = 40 d^−1^ (lower limit, longer half-life)). Next we scaled up *k*_*form*_ and *k*_*break*_ by the same amount, which yields the same percentage of T cells in synapses, However, the synapse half-life is shorter for the scaled-up *k*_*break*_ relative to the nominal value, thus, peak trimers-per-synapse reduce for increased *k*_*break*_ values (Figure S7A). Importantly, if we keep *k*_*form*_ constant but increase *k*_*break*_, the trimers-per-synapse magnitudes show the same quantitative behavior (Figure S7B, full transport model plots). Only *k*_*break*_ matters in this context.

**Figure 5:**
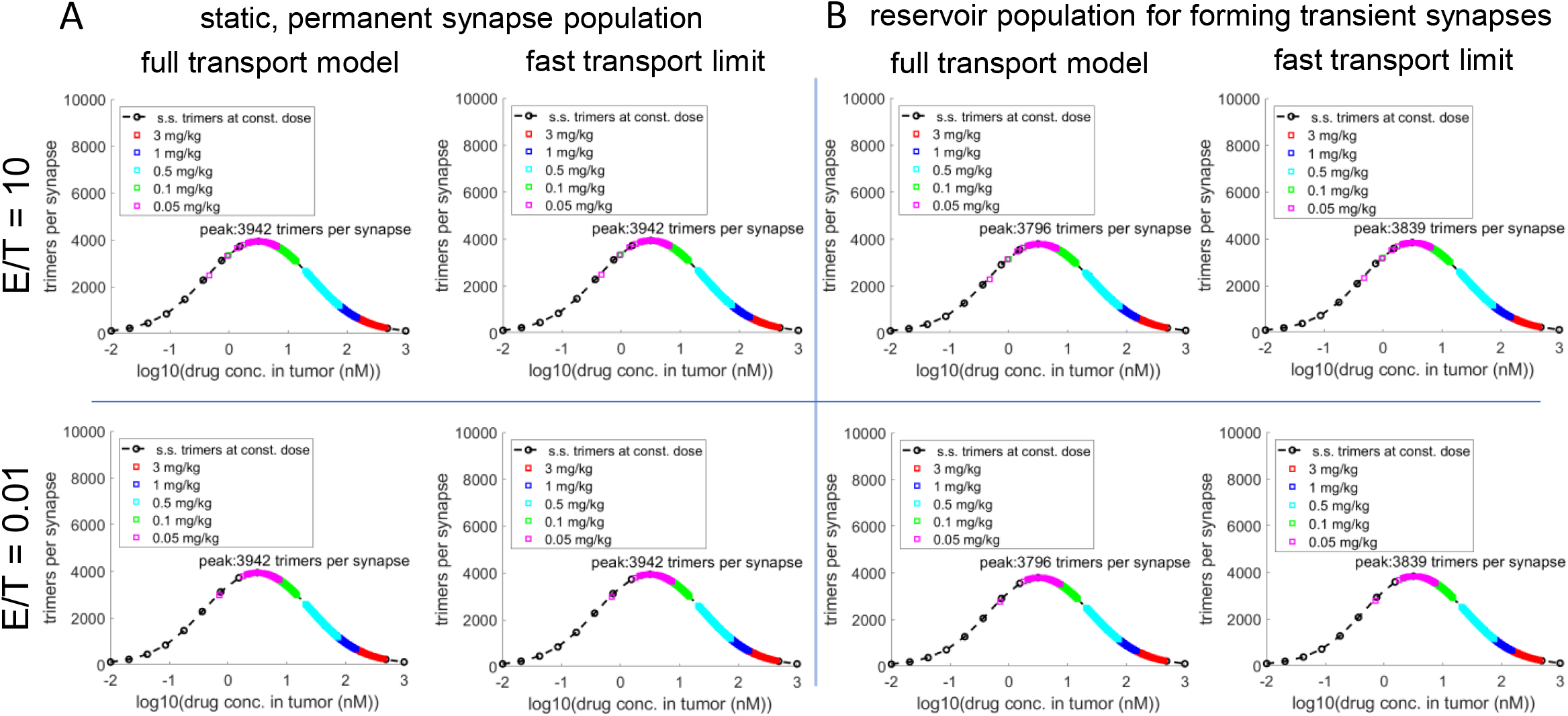
Comparison between trimer formation rate (full transport model) vs. trimer formation rate based on fast transport limit assumption (Eq. s3). A) Results for population of permanent synapses for the full transport model (left plot) and the fast transport limit model (right plot). B) Results for transient synapse population for the full transport model (left plot) and the fast transport limit model (right plot) where the synapse formation rate is *k*_*form*_ = 2 *×* 10^−11^ cells^−1^ L d^−1^, and the synapse break rate is *k*_*break*_ = 40 d^−1^. For each bell plot: colored diamonds represent dose-dependent, instantaneous values of trimers per synapse vs. dose-dependent, instantaneous values of tumor bsAb concentration (taken from the dose-dependent, time-dependent trimers per synapse plot in Figure S6). Black dashed-line with circles represents steady-state trimers per synapse vs. steady-state tumor bsAb concentration. For all cases, Γ = 200, and *KD*_*iaa*_ = 10 nM and *KD*_*taa*_ = 1 nM for the bsAb, and TAA expression levels were 10,000 receptors per cell. For the full transport model, the transport rate leaving the synapse is 10,000 d^−1^.

For the systems of equations used in this paper (permanent synapse model and transient synapse model), we refer the reader to Section S3.

### Application of the single-synapse model to simulating tumor growth inhibition

The invariance of the steady-state number of trimers per synapse as the E:T ratio is varied for a given bsAb concentration is a physical property of the single-synapse model as we highlighted above. It is how we would expect the biology to behave. We have shown that prior trimer models that don’t explicitly model synapses fail this test. However, that does not mean that these models are not useful for modeling tumor growth inhibition (TGI) experiments. Arguably, these models have done a reasonable job in this arena, e.g., Betts et al (2019) [10]. Part of the reason for their success may be that downstream of the trimer model is typically a target cell killing function that can compensate for the error in the trimer function which helps provide more flexibility in fitting. The killing function we employ for this example is the nonlinear hill function 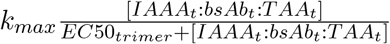 used in [10].

In Figure 6 we present results for two mouse TGI models, each with a different trimer model. The trimer models are the transient synapse model discussed above with the full trimer/transport model and the second model is from Betts et al (2019) [10]. We ran a simple numerical example for these tumor models where other than the type of trimer model and its rate constants, the remaining aspects of the two models are identical with the same rate constants and values. This includes the T-cell population growth/decay model and the tumor growth/inhibition model (see the transient synapse model in Section S3 for the mathematical form of these models). One exception is that the *EC*50_*trimer*_ value of the tumor inhibition model is different between our syanpse-based TGI model and the Betts TGI model.

**Figure 6:**
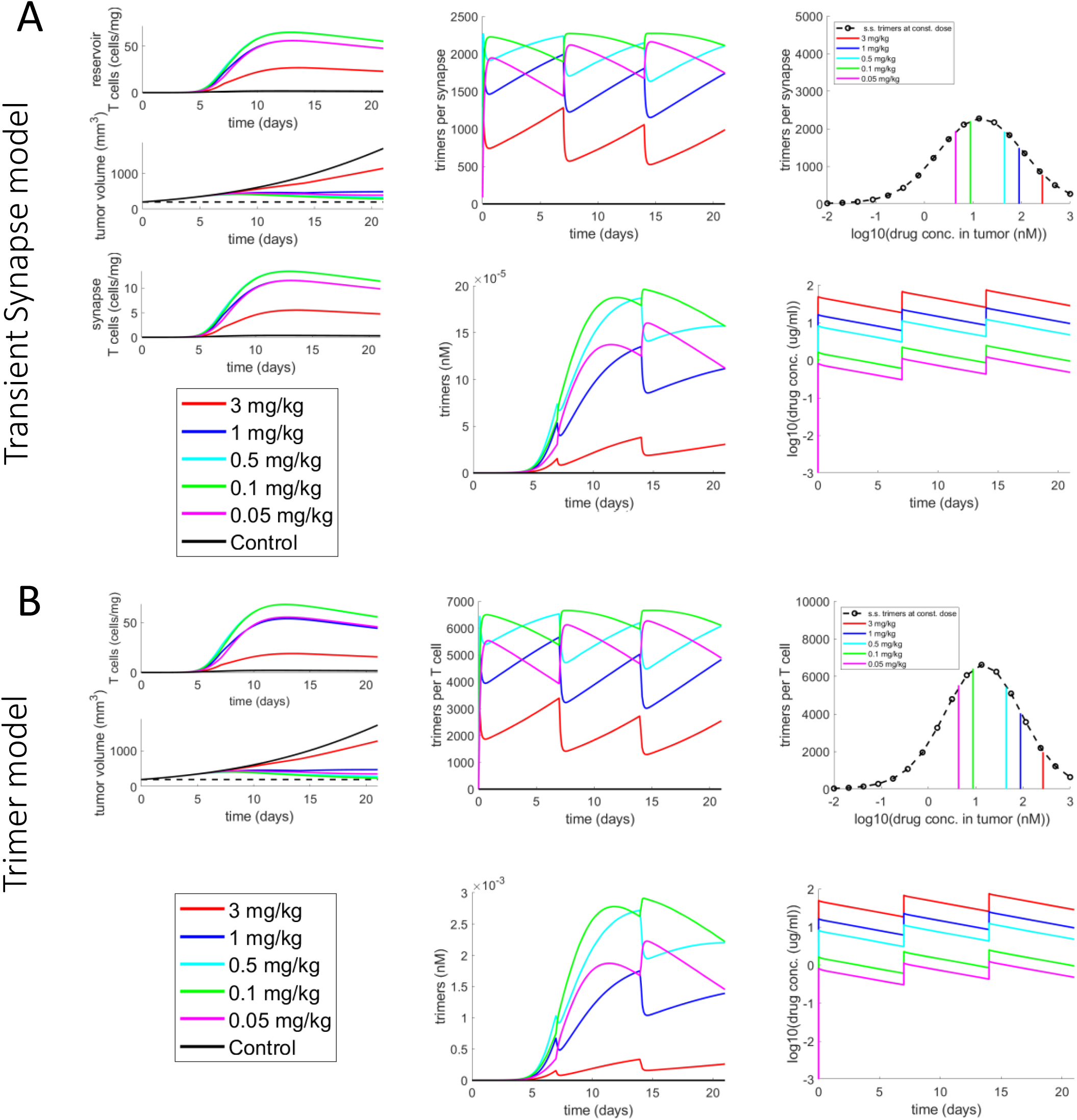
TGI models with dynamic T cell populations. A) Transient synapse model: reservoir population of T cells that drive a small synapse population that constantly form transient synapses with full transport model and Γ = 500. B) Trimer model from [10]: Single T cell population. Left column: T cell population(s) and TGI. Middle column: time-dependent trimers per synapse (or trimers per T cell (Trimer model)), time-dependent total trimers (nM). Right column: trimers-per-synapse (or trimers per T cell (Trimer model)) vs. tumor bsAb concentration, PK. For bell curve: Vertical colored bars represent dose-dependent, time-averaged trimers per synapse (or trimers per T cell (Trimer model)) vs. dose-dependent, time-averaged tumor bsAb concentration. Black dashed-line with circles represents steady-state trimers per synapse (or trimers per T cell (Trimer model)) vs. steady-state tumor bsAb concentration. For both cases, *KD*_*iaa*_ = 64 nM and *KD*_*taa*_ = 3.22 nM for the bsAb, and TAA expression levels were 10,000 receptors per cell.

The top group of plots represent results from a tumor model that applies the transient synapse model where T cells from the T cell population are transiently forming synapses with target cells, physiologically closer to reality than the Betts model. In these simulations the percentage of T cells in synapses are on the order of 20 percent of the total T cell population in the tumor. Overall, the TGI is similar across the models. Here we adjusted some of the parameters by hand such that they show similar results.

For the synapse model (Figure 6A), one observes that the synapse population is dynamically changing, first rising then eventually falling over time, as is the total T cell population in the tumor. Mirroring this is the plot of total trimers, while the plot of trimers-per-synapse vs. time is invariant to the dynamically changing E:T ratio. Furthermore, their average trimers per synapse vs. dose maps well to the steady-state trimers per cell plot, as we observed in earlier examples.

For the transient synapse model, both transport rates for IAAs and TAAs in and out of the synapse and the synapse half-life (discussed above) are important for reaching a certain level of trimers per synapse given Γ. If transport is too slow the number of trimers formed will be lower (see Figure S8), both total trimers and trimers per synapse.

Given our synapse model’s ability to calculate trimers-per-synapse, we now have the ability to model nonlinear/thresholded trimers-per-synapse activation that may exist at the single-synapse level. This can then be multiplied by the trimer concentration killing function and ensure that not only are the number of trimers important, but that the trimers-per-synapse in each synapse is driving a high enough level of activity. With regards to implementation of the activation function, we use a hill function

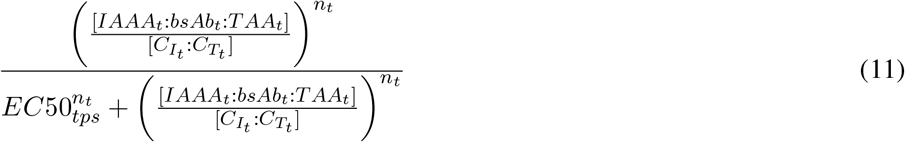

where the higher the *n*_t_ the more threshold-like the function is at 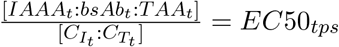. Applying this to the killing function yields a thresholded trimer killing function of the form

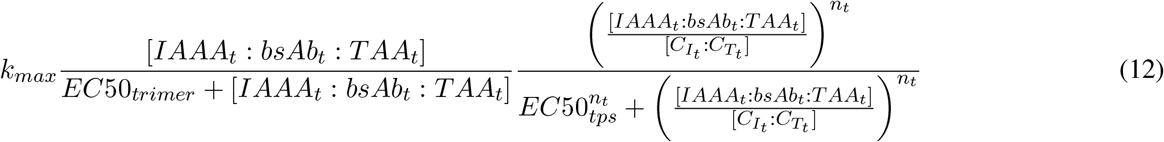

which has the same form and variables as the simple trimer killing function, but multiplied by trimer-per-synapse dependent activation function. While this approach may not always be needed for fitting data, it may be useful for fitting data that shows a threshold dependence and, thus, provides some extra flexibility. We compare the simple trimer killing function and the thresholded trimer killing function for some simple TGI examples with a constant E:T ratio (Figure 7A). Here for a given trimer killing function, we play with the trimers-per-synapse threshold (*EC*50_*tps*_) for the activation function and where *n*_*t*_ = 4. Furthermore, the Betts model cannot apply thresholding accurately (or any non-synapse model for that matter) since it doesn’t model synapses. For increasing *EC*50_*tps*_ threshold values of 20, 2000, and 4000, the dose-dependent TGI behavior decreases as would be expected.

**Figure 7:**
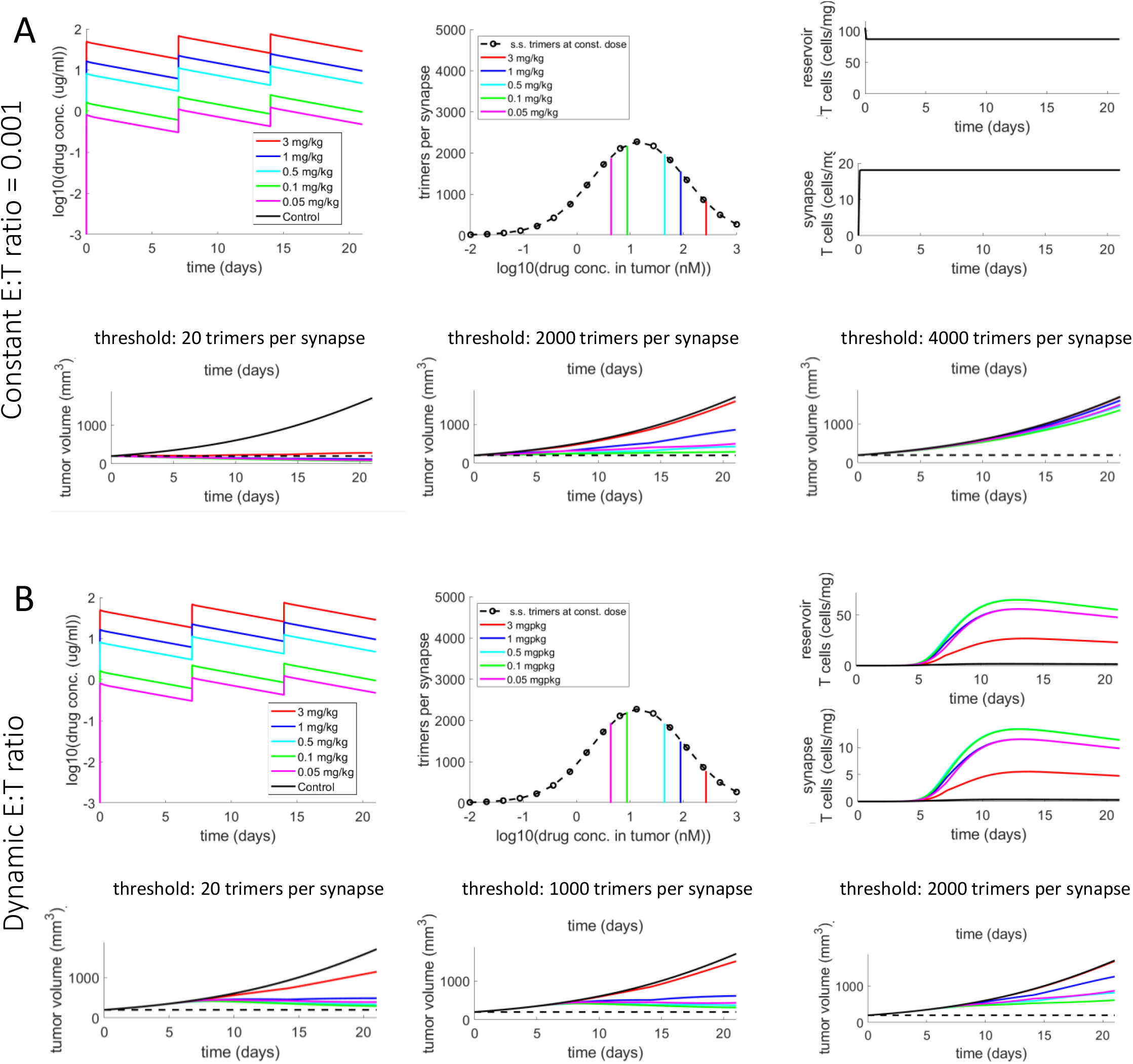
TGI for different trimers-per-synapse activation thresholds. For A and B: Top panels: left: dose-dependent PK, middle: trimers per synapse, right: T cell dynamics. Bottom panels: TGI for different trimers-per-synapse activation thresholds. A) Constant E:T ratio = 0.001 with threshold *EC*50_*tps*_ values of 20 trimers per synapse, 2000 trimers per synapse, and 4000 trimers per synapse, respectively, and *n*_*t*_ = 4. B) Dynamic E:T ratio with threshold *EC*50_*tps*_ values of 20 trimers per synapse, 1000 trimers per synapse, and 2000 trimers per synapse, respectively, and *n*_*t*_ = 4. For all simulations, Γ = 500.

We next applied the thresholded trimer killing function to the dynamic E:T ratio case from Figure 6A. For this TGI model, everything upstream of the TGI are quantitatively the same, e.g. dose-dependent trimers, trimers per synapse, etc. (compare (A) and (B) in Figure S9). With regards to TGI, for increasing *EC*50_*tps*_ values of 20, 1000, and 2000, this leads to a decreasing shift in the dose-dependent TGI (Figure 7B). To reiterate, the overall goal of this approach is to capture more complex dose-dependent TGI signatures with this model that simpler killing functions cannot.

## Discussion

In this work, we have presented a single-synapse model that conserves certain properties such as the number of trimers per synapse for a given bsAb concentration, even as the E:T ratio is changed. This property is not captured in the formulation of other ODE trimer models in the literature, and represents a fundamental improvement in ODE trimer models. While agent based models naturally conserve this since each synapse complex is modeled independently, they tend to be much more computationally intensive and complicated [18]. Furthermore, we introduce new capabilities such as modeling transient synapse formations where only a certain portion of T cells from the population are in synapses at any given time. Overall, this allows our model to be coupled to more complex immune cell population interactions if necessary. Our synapse model is modestly more complex than prior trimer models, but allows for much greater freedom as well as proven calculations that are more consistent across E:T ratios at a given bsAb concentration.

We wanted to explore ways of possibly simplifying (coarse graining) our synapse model. If we enforce extremely fast timescales of species redistribution, a given non-trimer species will have the same concentration inside and outside the synapse, when there are trimers (presence of bsAb) or not (no bsAb). Given this concentration constraint, we do not have to explicitly model transport in and out of the synapse, which will yield a slightly different form of the reaction rate for the trimer (Eq. s3). See Section S1 for the derivation of Eq. (s3).

For permanent synapses, the modeling approach using Eq. (s3) agrees well with our results for our full transport model (Figure 5A, compare full transport model vs. fast transport limit). This fast transport limit model represents an upper bound on steady-state trimer formation for the cases presented. However, for the finite synapse case in Figure 5B (compare full transport model vs. fast transport limit), simulations using Eq. (s3) have higher peak trimers per synapse than that for the full model with transport. As one increases the value of *k*_*break*_ to *k*_*break*_ = 320 d^−1^ and *k*_*break*_ = 640 d^−1^ the peak trimers per synapse decreases more strongly in the full transport model than the fast-transport limit model (Figure S7B, compare full transport model plots to fast transport limit model plots). This is because impact of the timescale of transport on peak trimers per synapse becomes more important as the synapse half life decreases (*k*_*break*_ gets larger). As expected, at a given steady-state dose, the trimers per synapse will be invariant across E:T ratios for a particular *k*_*break*_ value (Figure S7). The values of *k*_*break*_ used are all within observed half-lifes of synapses [29, 30]. Because of the greater flexibility in better modeling the effects of different *k*_*break*_ and transport rates, our full model with transport is definitely more general.

The next step for our single-synapse model for bispecific immune cell engagers is to test how well it is suited to be a translation/predicive tool for estimating FIH doses and efficacy in the clinic. The plan is to test it across different internal and published bispecific-oncology drug programs with available data sets.

For the trimer bimolecular reaction rate derived above, i.e., Eqs (3) and (4), we assume an nM system where trimers are in nM and 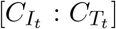 is in synapse complexes per liter. We can easily adapt the equations to other equivalent systems, especially those at constant volume. For an nmol system that is solving for trimers in nmol, in Eqs (3) and (4), one would just define 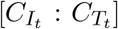 as the number of synapse complexes in the tumor interstitial volume and the derivations would look the same and simulation results would be identical when converted to nM. However, Γ would be different between both systems in both value and dimension, but related by a conversion factor since they must yield the same results.

We focused mainly on a bispecific antibody with monovalency for both the TAA arm and the IAA arm (1+1 format). There are other bispecific formats such as bivalency for the TAA arm and monovalency for the IAA arm (2+1 format), an example being the drug Xaluritamig (AMG509) which has shown promise in the clinic for prostate cancer patients [33, 34, 35]. And there are also trispecific antibody formats and beyond. Our method is capable of modeling these various drug types.

For the work presented here, we are only modeling single-synapse complexes, but in reality, multi-synapse complexes have been observed *in vitro* [18] in significant numbers for up to *N* of 3 synapses per complex, although the single-synapse complex is still the dominant population from these observations. In vivo, E:T ratios tend to be in favor of tumor cells [36]. Furthermore, T cells can only engage in one active synapse at a time due to having only one centriole [37, 38, 39, 40]. Hence, in vivo, single-synapse modeling is likely most appropriate. Still, multi-synapse complexes may still occur *in vivo* when the E:T ratio is in favor of T cells (likely due to significant T cell infiltration to the site of action), where more than one T cell is engaged with a single tumor cell. For each of the observed multi-synapse complex types, we do expect the trimers per synapse to be invariant across E:T ratios at a given bsAb concentration, but likely different from the others including the single-synapse complex. This is because each multi-synapse complex types is unique in how the TAA on the tumor cell(s) and IAA on the immune cell(s) are shared across the synapses at a given bsAb concentration. For instance, the more synapses on a tumor cell the less TAA available for trimers in each synapse. A current focus of research is to extend our model to include multi-synapse complexes and test head-to-head the fitting, prediction, and translational capabilities of the single-synapse and multi-synapse models.

The transport model only involved two compartments (synapse and non-synapse regions). More non-synapse compartments could be used to better model the diffusion-driven transport. This could be tested in future models.

Because our synapse model is general, we can easily apply the approach to other immune cell therapies such as Chimeric Antigen Receptor T cells (CAR-T) as well as general immune cell/target cell interactions. See S2 for some simple demonstrations of the approach.

For this paper, we purposefully modeled constant total receptors per cell with no synthesis nor internalization to demonstrate the mathematical properties of trimers per synapse across E:T ratios at a given bsAb concentration. In general, receptor synthesis, internalization (and potentially recycling) can be modeled. The invariance properties discussed in this paper should hold, but the magnitude of the trimers per synapse at a given concentration will likely take on a smaller value since internalization will break/remove trimers.

For the types of expansions of the model discussed above, it is important to practice caution when applying them such as when there are data sets to support them or if they are required for translation. Barring that, the model we presented in this work serves as both a useful model and a foundation for future models.

---

**Figure S1:**
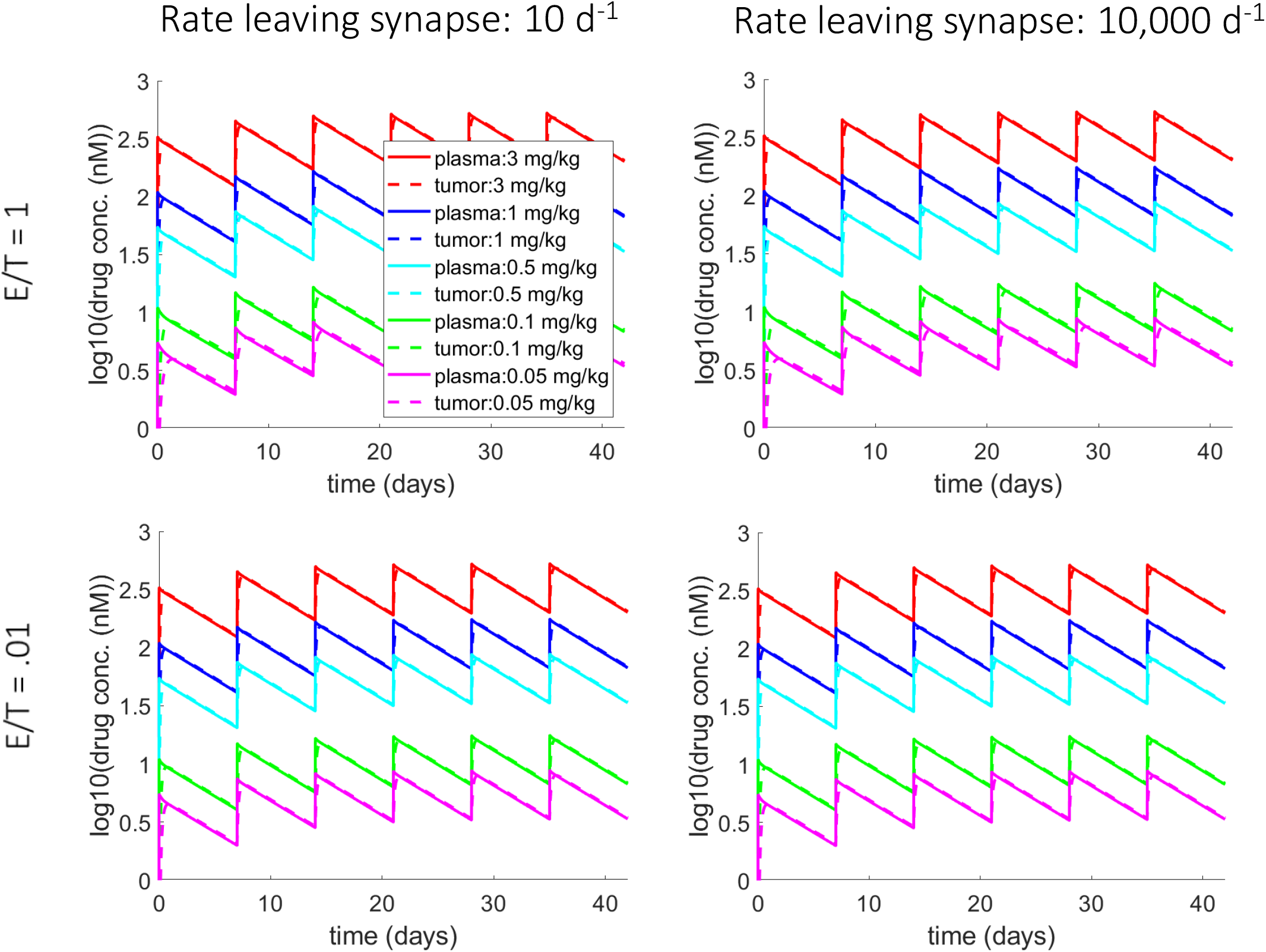
Dose-dependent plasma and tumor PK for the trimers per synapse calculations from Figure 2. For these simulations, the bsAb has a *KD*_*iaa*_ = 10 and *KD*_*taa*_ = 1. For the synapse transport rate leaving the synapse, simulations were run for values of 10 and 10000 d^−1^, E:T ratios of 1 and 0.01, doses of 0 mg/kg (control), 0.05 mg/kg, 0.1 mg/kg, 0.5 mg/kg, 1 mg/kg and 3 mg/kg, and Γ = 200. For each synapse transport rate and E:T ratio, the dose-dependent plasma and tumor bsAb concentrations over time are plotted.

**Figure S2:**
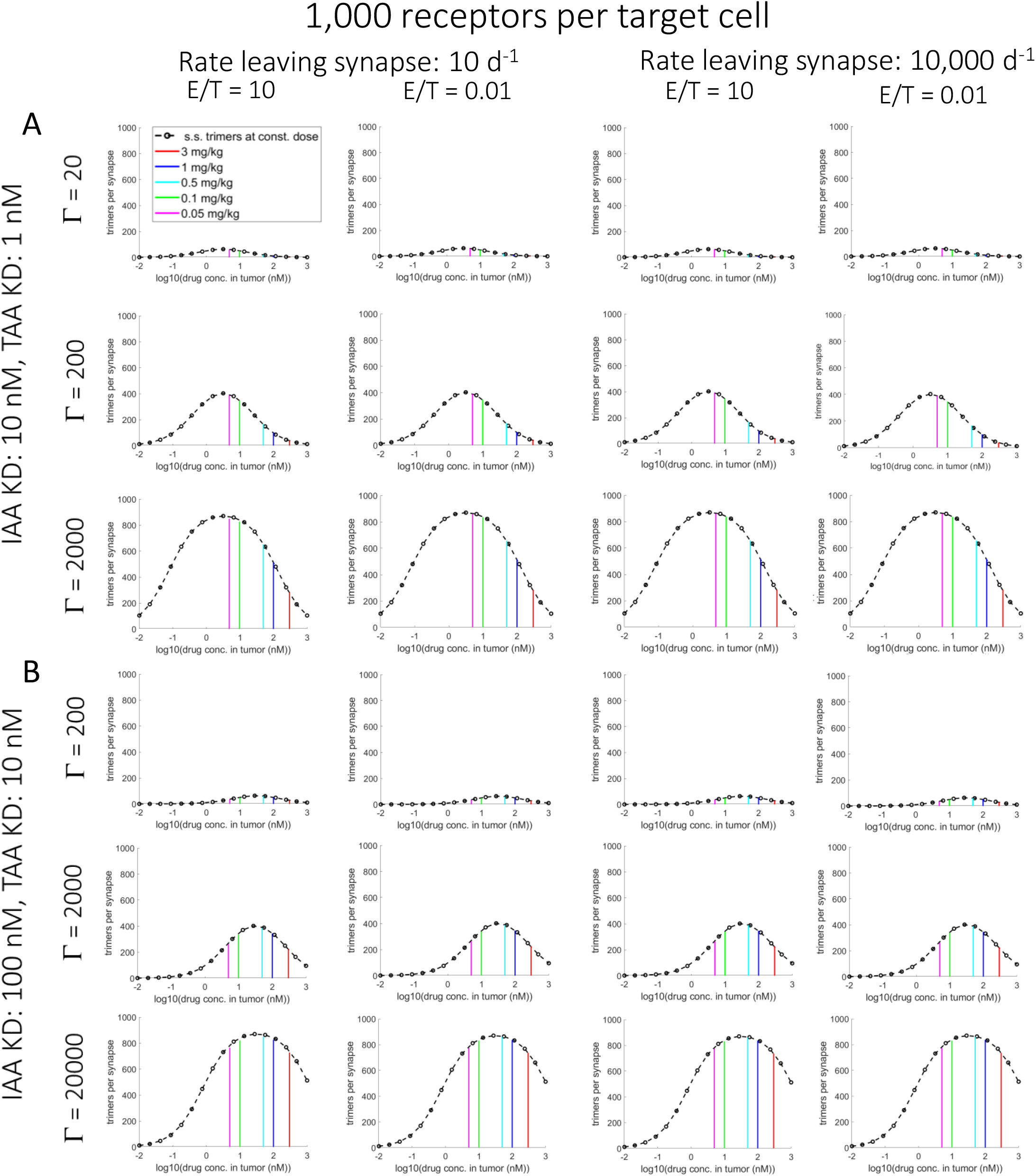
Synapse model results of trimers per synapse vs. bsAb tumor concentration for 1,000 receptors per target cell are invariant over E:T ratios. Simulations were run for different bsAb paired KD values: (A) *KD*_*iaa*_ = 10 and *KD*_*taa*_ = 1 for Γ values of 20, 200, and 2000. (B) *KD*_*iaa*_ = 100 and *KD*_*taa*_ = 10 for Γ values of 200, 2000, and 20000. For (A) and (B) simulations were run for transport rates leaving the synapse of 10 d^−1^ and 10000 d^−1^, E:T ratios of 10 and 0.01, and doses of 0.05 mg/kg, 0.1 mg/kg, 0.5 mg/kg, 1 mg/kg, and 3 mg/kg. For all plots: Vertical colored bars: dose-dependent, time-averaged trimers per synapse vs. dose-dependent, time-averaged tumor bsAb concentration. Black dashed-line with circles: steady-state trimers per synapse vs. steady-state tumor bsAb concentration.

**Figure S3:**
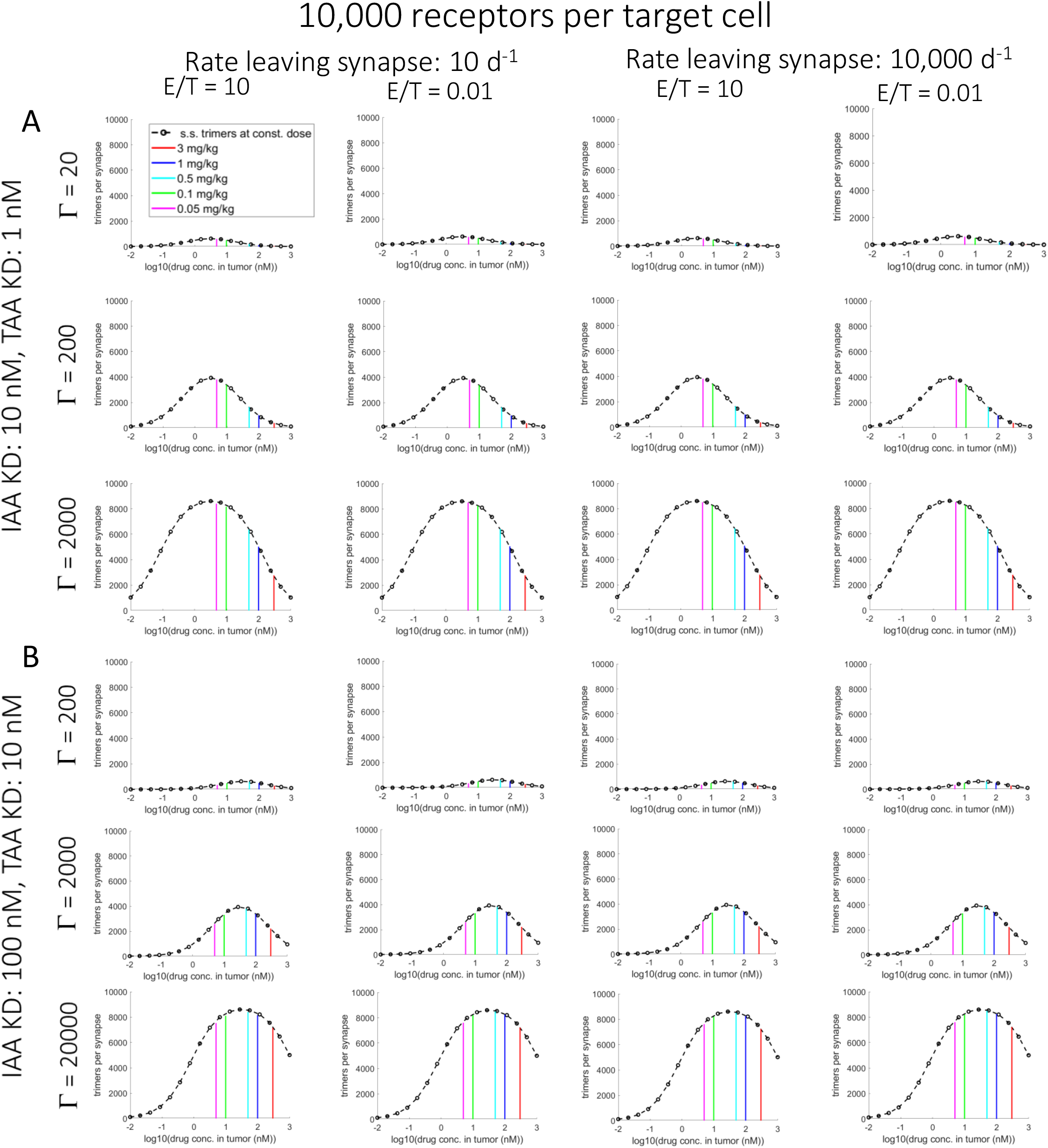
Synapse model results of trimers per synapse vs. bsAb tumor concentration for 10,000 receptors per target cell are invariant over E:T ratios. Simulations were run for different bsAb paired KD values: (A) *KD*_*iaa*_ = 10 and *KD*_*taa*_ = 1 for Γ values of 20, 200, and 2000. (B) *KD*_*iaa*_ = 100 and *KD*_*taa*_ = 10 for Γ values of 200, 2000, and 20000. For (A) and (B) simulations were run for transport rates leaving the synapse of 10 d^−1^ and 10000 d^−1^, E:T ratios of 10 and 0.01, and doses of 0.05 mg/kg, 0.1 mg/kg, 0.5 mg/kg, 1 mg/kg, and 3 mg/kg. For all plots: Vertical colored bars: dose-dependent, time-averaged trimers per synapse vs. dose-dependent, time-averaged tumor bsAb concentration. Black dashed-line with circles: steady-state trimers per synapse vs. steady-state tumor bsAb concentration.

**Figure S4:**
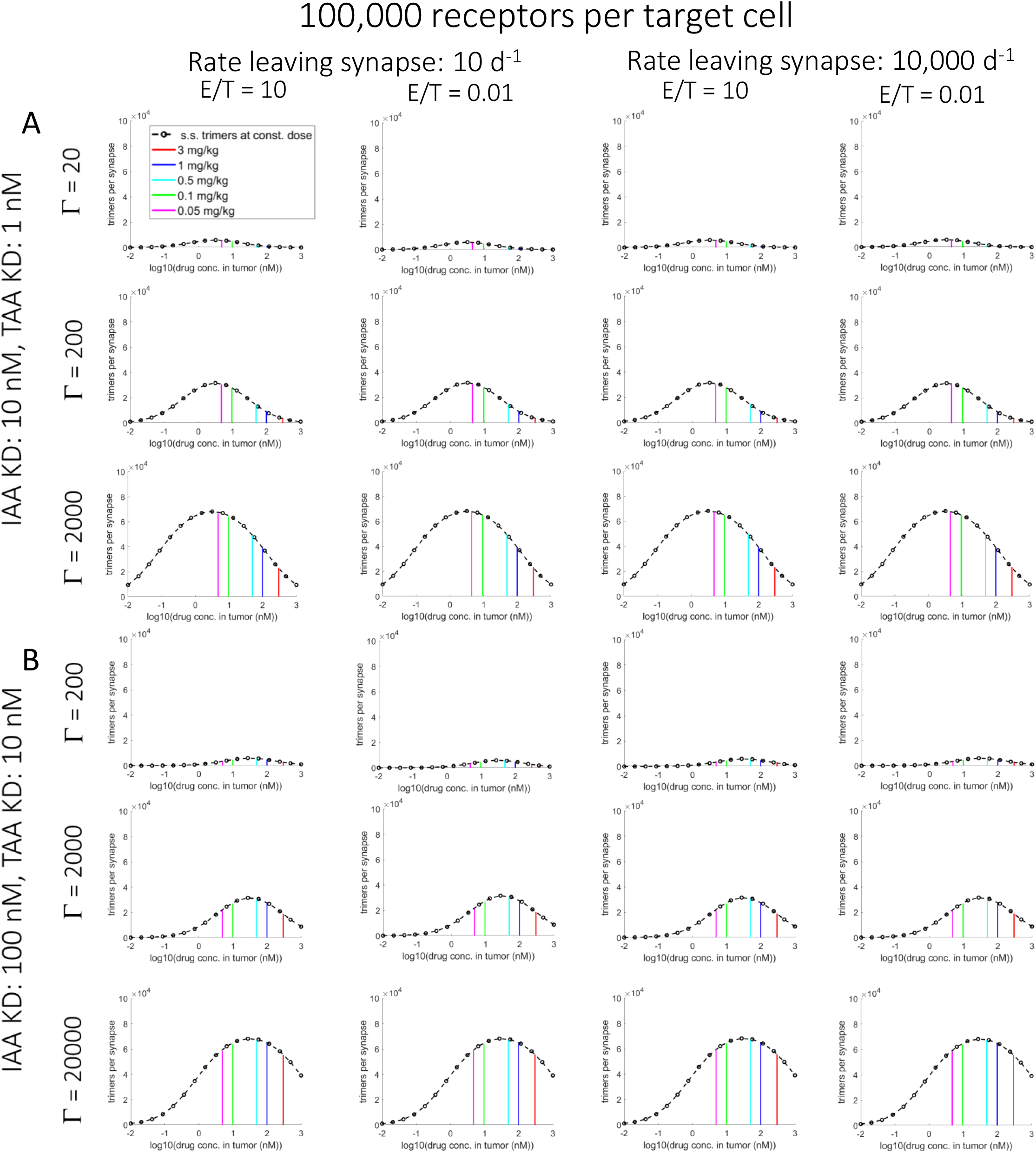
Synapse model results of trimers per synapse vs. bsAb tumor concentration for 100,000 receptors per target cell are invariant over E:T ratios. Simulations were run for different bsAb paired KD values: (A) *KD*_*iaa*_ = 10 and *KD*_*taa*_ = 1 for Γ values of 20, 200, and 2000. (B) *KD*_*iaa*_ = 100 and *KD*_*taa*_ = 10 for Γ values of 200, 2000, and 20000. For (A) and (B) simulations were run for transport rates leaving the synapse of 10 d^−1^ and 10000 d^−1^, E:T ratios of 10 and 0.01, and doses of 0.05 mg/kg, 0.1 mg/kg, 0.5 mg/kg, 1 mg/kg, and 3 mg/kg. For all plots: Vertical colored bars: dose-dependent, time-averaged trimers per synapse vs. dose-dependent, time-averaged tumor bsAb concentration. Black dashed-line with circles: steady-state trimers per synapse vs. steady-state tumor bsAb concentration.

**Figure S5:**
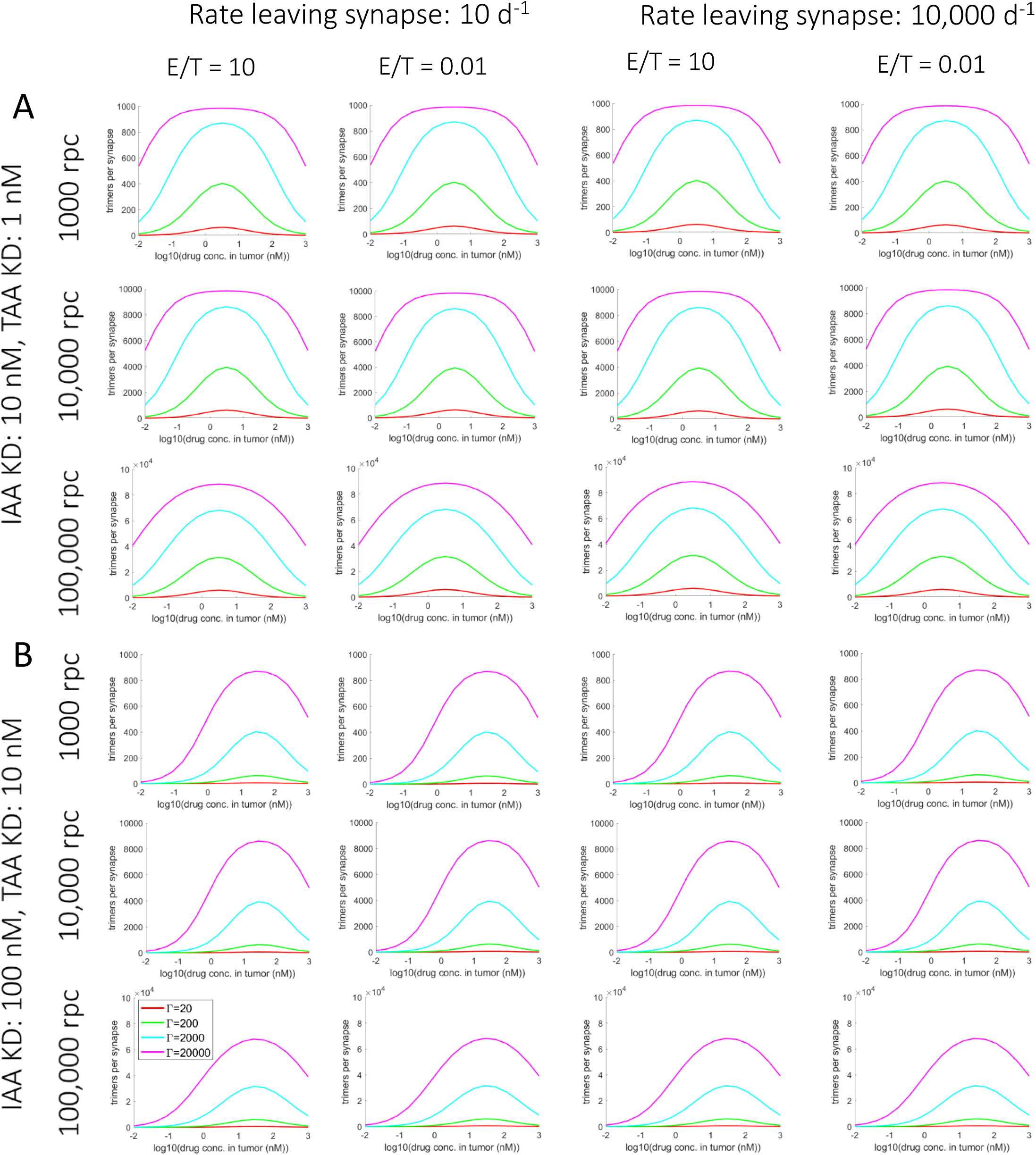
Summary of synapse model results of steady-state trimers per synapse vs. steady-state tumor bsAb concentration that are invariant over E:T ratios and a large range of transport timescales. Simulations were run for different bsAb paired KD values: (A) *KD*_*iaa*_ = 10 nM and *KD*_*taa*_ = 1 nM. (B) *KD*_*iaa*_ = 100 nM and *KD*_*taa*_ = 10 nM. For (A) and (B), simulations were run for different TAA expression levels (1000, 10,000, 100,000 receptors per cell). For these different combinations, simulations were run for different transport rates exiting the synapse of 10 d^−1^ and 10000 d^−1^, E:T ratios of 10 and 0.01, and Γ values of 20 (red), 200 (green), 2000 (cyan), 20000 (magenta).

**Figure S6:**
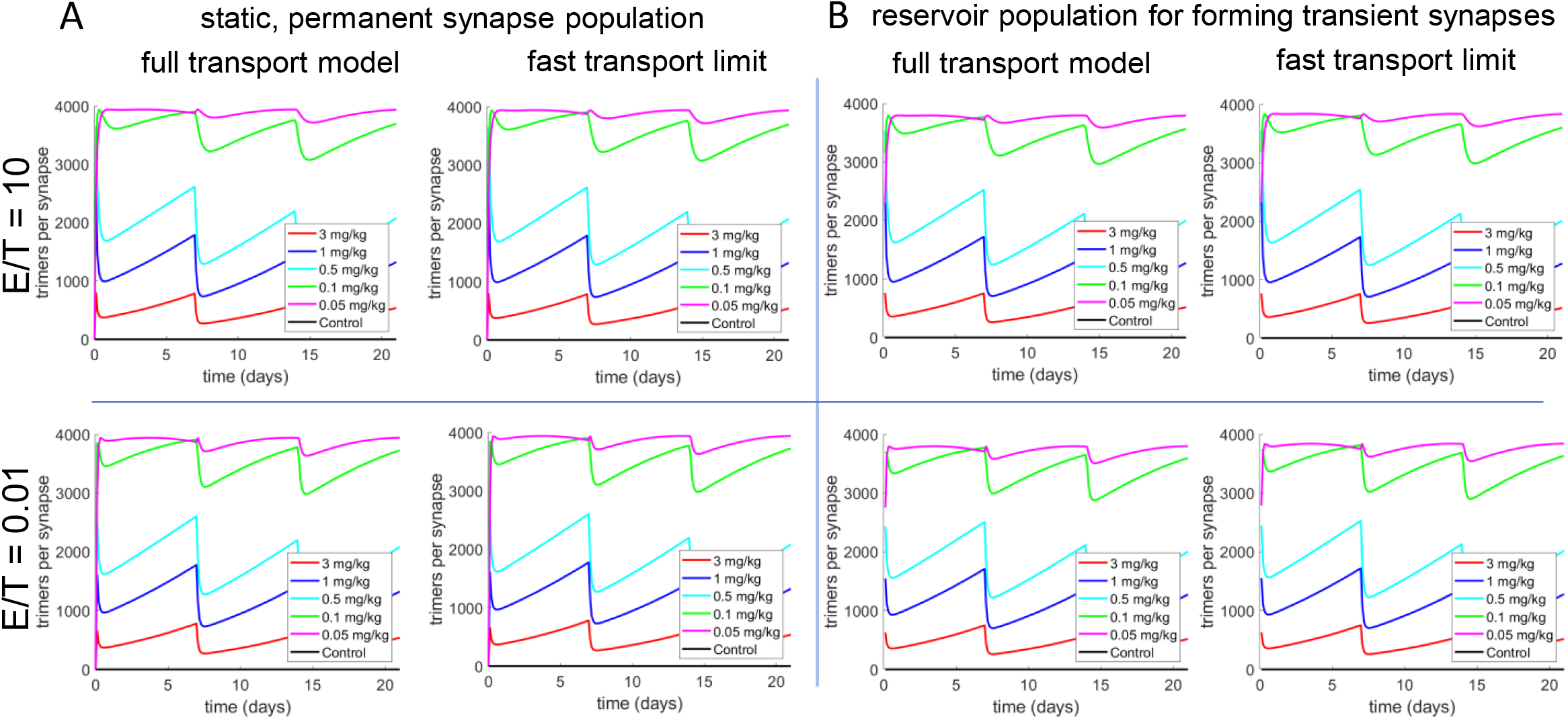
Time-dependent trimers-per-syanpse comparison between trimer formation rate (full transport model) vs. trimer formation rate based on fast transport limit assumption (Eq. s3). A) Results for population of permanent synapses for the full transport model (left plot) and the fast transport limit model (right plot). B) Results for transient synapse population for the full transport model (left plot) and the fast transport limit model (right plot) where the synapse formation rate is *k*_*form*_ = 2 *×* 10^−11^ cells^−1^ L d^−1^, and the synapse break rate is *k*_*break*_ = 40 d^−1^. For each case above, the dose-dependent, time-dependent trimers per synapse plots (dose-dependent tumor PK plot not shown)) were simulated for E:T ratios of 10 and 0.01, respectively. For all cases, Γ = 200, and *KD*_*iaa*_ = 10 nM and *KD*_*taa*_ = 1 nM for the bsAb, and TAA expression levels were 10,000 receptors per cell. For the full transport model, the transport rate leaving the synapse is 10,000 d^−1^.

**Figure S7:**
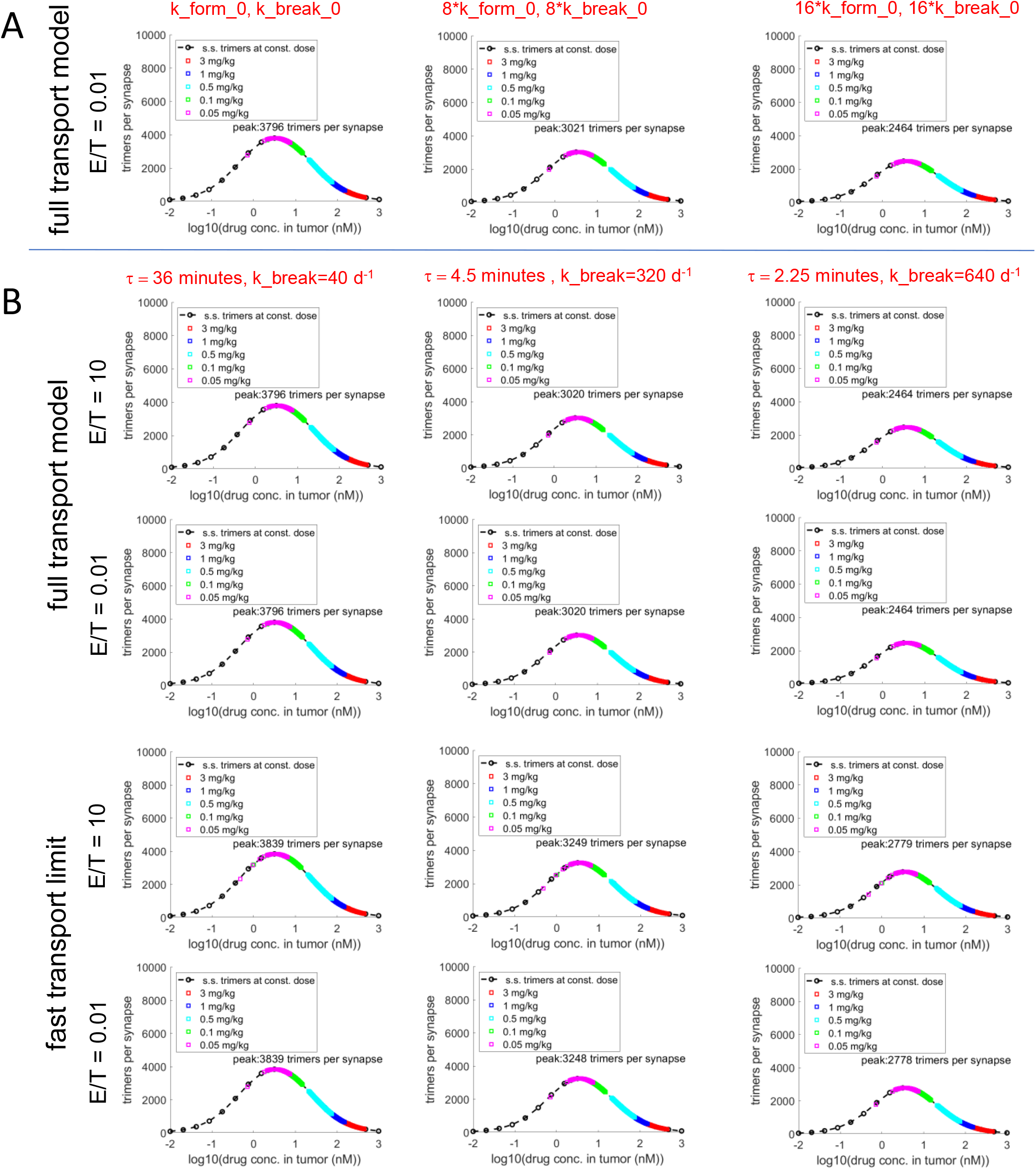
Magnitude of synapse break rates affect peak level of trimers per synapse for the transient synapse model. Maintaining a constant synapse populations size with proportional increases in both *k*_*form*_ and *k*_*break*_. Here 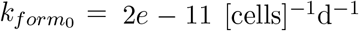 and 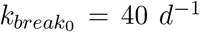, i.e. the nominal values. Percentage of T cells in synapses is 20.6% across cases. (B) Simulations for the nominal value of *k*_*form*_ and increasing values of *k*_*break*_ for the full tranport model and the fast transport limit model (Eq. s3). Simulated values of *k*_*break*_ were: 40 *d*^−1^ (left column), 320 *d*^−1^ (middle column), and 640 *d*^−1^ (right column). For each model in (B), top row: E:T ratio of 10, bottom row: E:T ratio of 0.01. For all cases, Γ = 200, and *KD*_*iaa*_ = 10 nM and *KD*_*taa*_ = 1 nM for the bsAb, and TAA expression levels were 10,000 receptors per cell.

**Figure S8:**
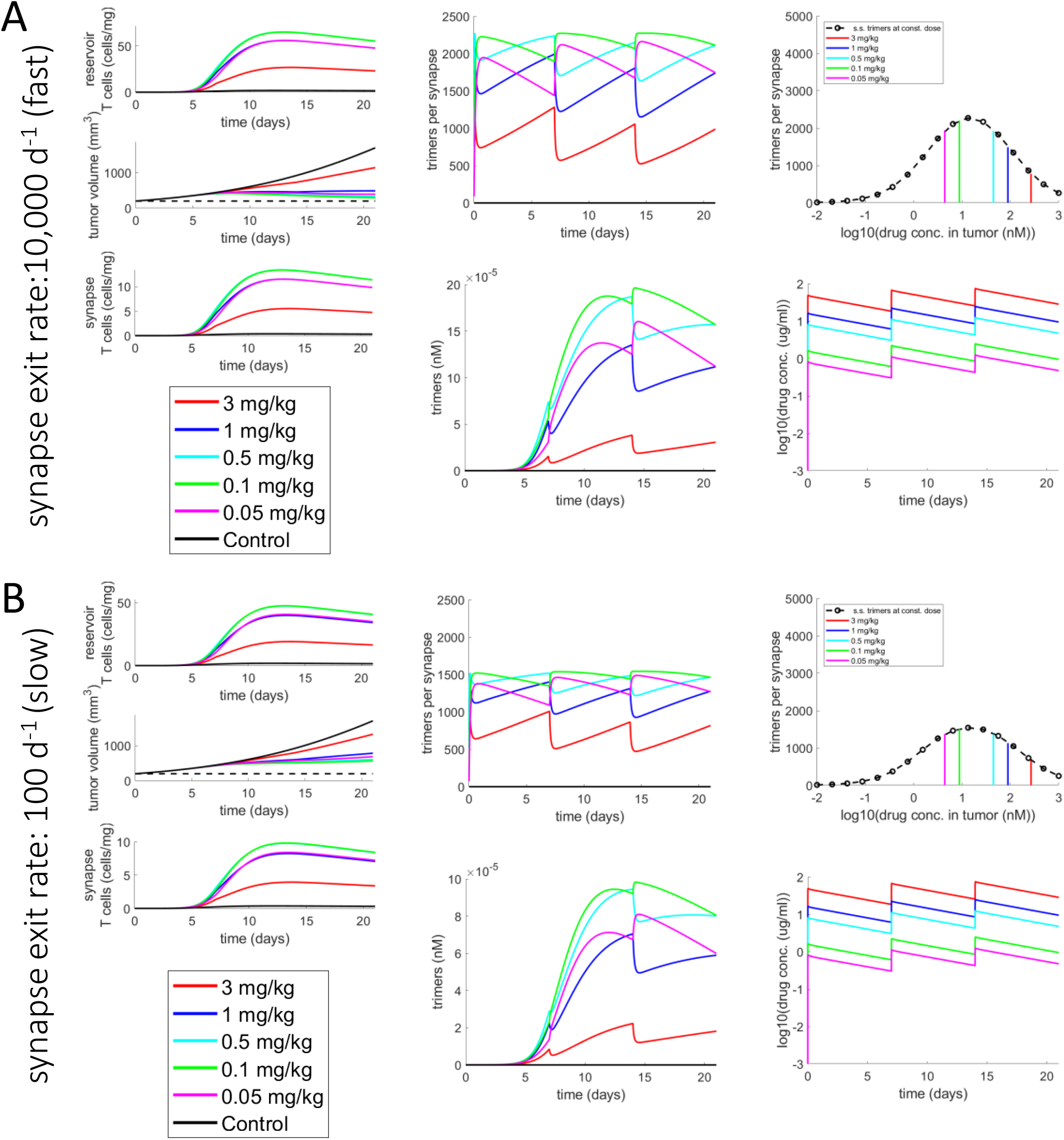
Magnitude of transport rates in and out of synapse affects peak level of trimers per synapse for the transient synapse model. (A) Transient synapse model from Figure 6A where the transport rate exiting the synapse is 10,000 d^−1^. Same as (A) but where the transport rate exiting the synapse exit rate is 100 d^−1^ (slow). For both cases, Γ = 500.

**Figure S9:**
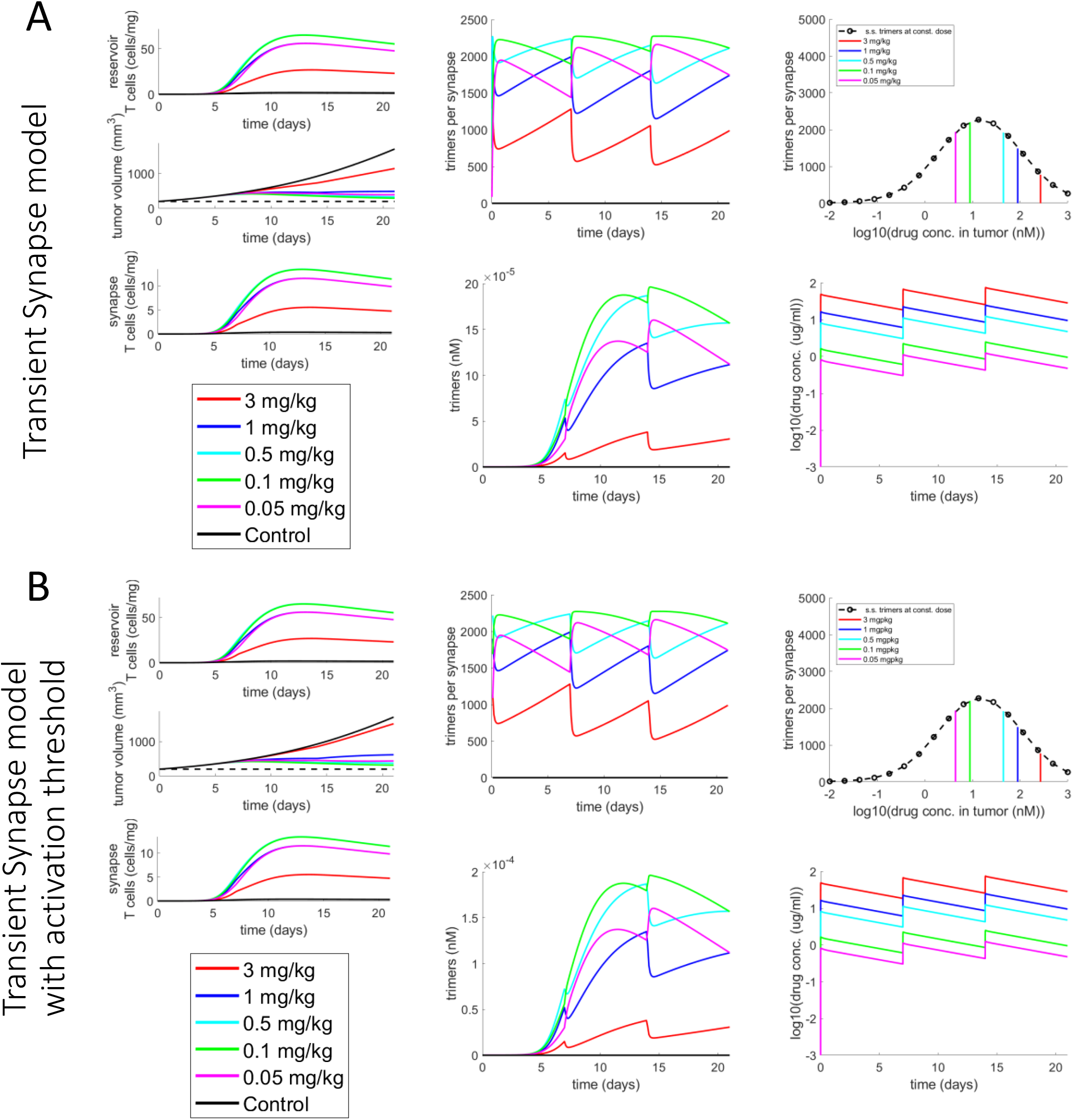
Including the trimers-per-synapse activation function to a TGI model with a dynamic T-cell population. (A) Transient synapse model from Figure 6A which uses the simple trimer killing function. (B) Same model as (A) but with thresholded trimer killing function where the trimers-per-synapse activation function has *EC*50_*tps*_ = 1000 and *n*_*t*_ = 4. For both cases, Γ = 500.

**Figure S10:**
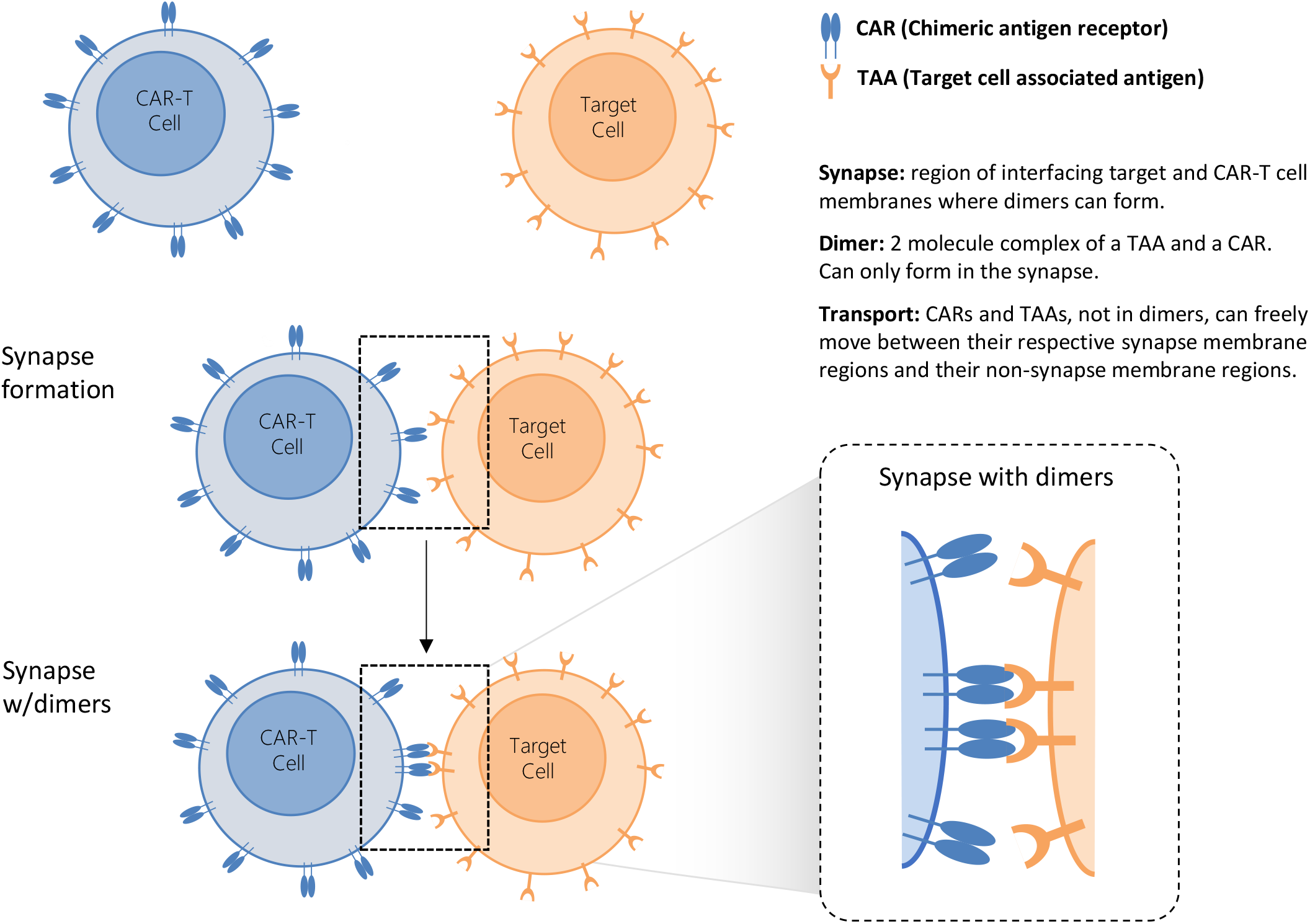
Formation of synapse with dimer connections between a CAR-T cell and target cell. Illustration shows a CAR-T cell and target cell (top two cells) that come together to form a synapse (middle two cells), i.e,, an interface of their respective cell membranes. Within the synapse dimers, (CAR-TAA) can form between the CAR-T cell and target cell (bottom two cells). With sufficient numbers of dimers, this can lead to CAR-T cell killing of target cells.

**Figure S11:**
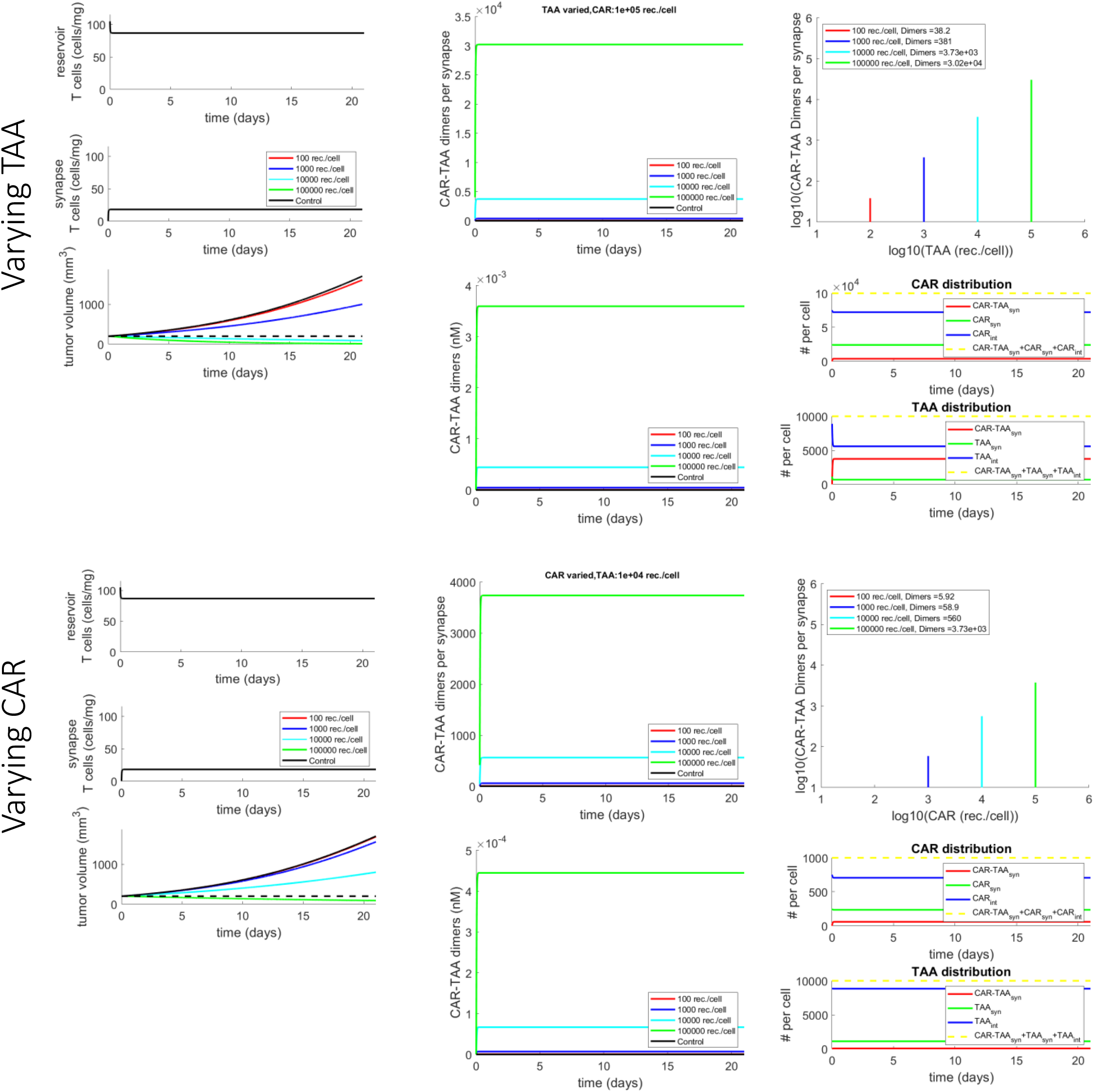
Simple CAR-T model at constant E:T ratio produces different levels of TGI when varying the TAA expression or CAR expression. These simulations modeled a reservoir population of CAR-T cells that drive a small synapse population that constantly form transient synapses. Top panels: varying TAA expression. Bottom panels: varying CAR expression. Left column: CAR-T cell dynamics and TGI. Middle column: dimer dynamics in dimers per synapse (top plot) and nM (bottom plot). Right column: dimers per synapse at the end point in time for different expression levels (top plot), receptor transport dynamics between synapse and non-synapse regions for a particular case (bottom plot) where TAA and CAR expression levels per cell are the numbers at the top of their respective y-axis. Here the avidity factor Γ = 5 with an E:T ratio of 0.001.

**Figure S12:**
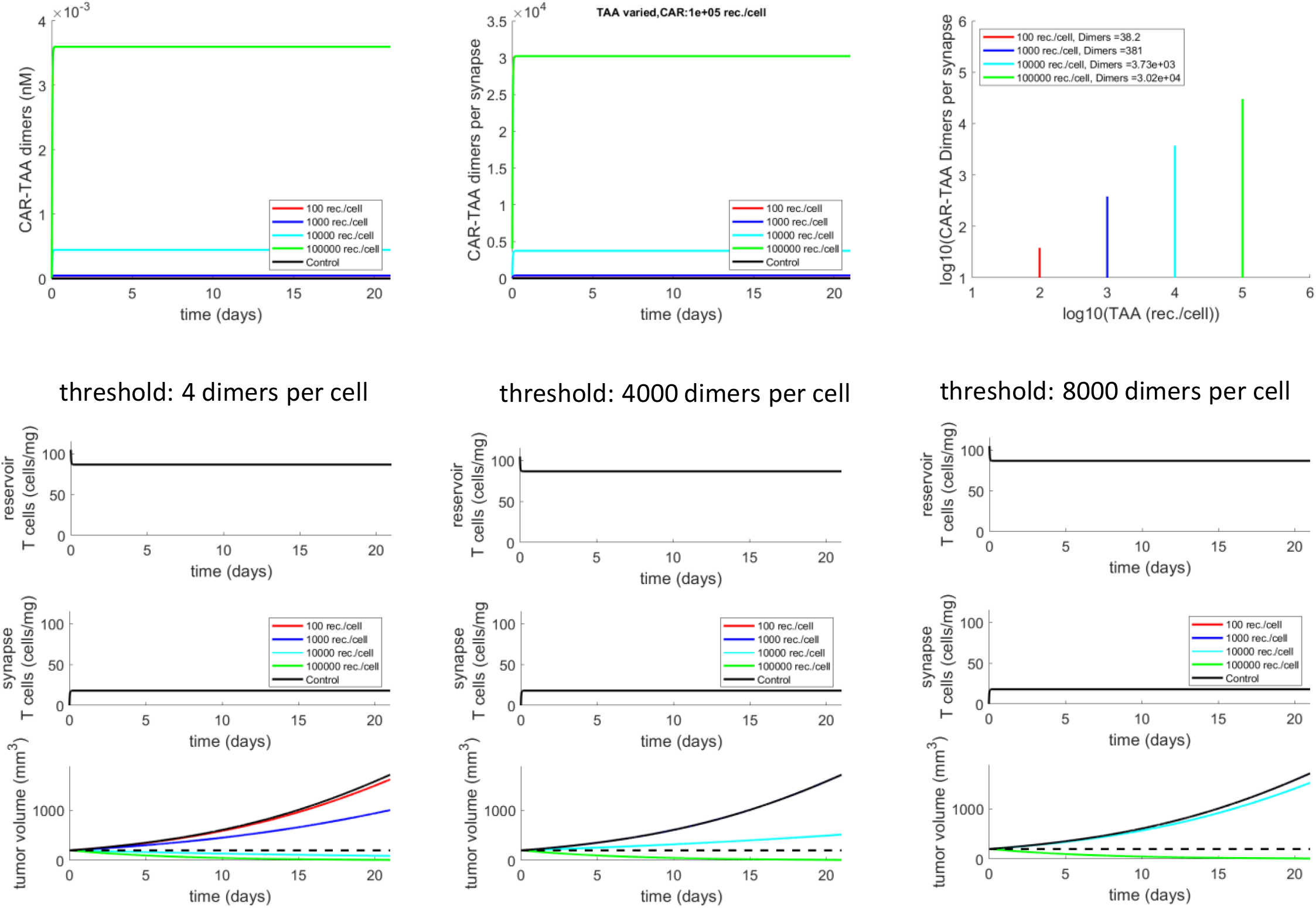
TGI for CAR-T model at constant E:T ratio for different dimers-per-synapse activation thresholds at a constant CAR expression level but varying TAA expression. Top panel: left and middle: dimer dynamics for different TAA expression levels, right: dimers per synapse at the end point in time for different TAA expression levels. Bottom panel: CAR-T cell dynamics and TGI for different dimers-per-synapse activation thresholds of 4, 2000, and 4000 dimers per synapse, respectively. Here the avidity factor is Γ = 5 with an E:T ratio of 0.001.

**Figure S13:**
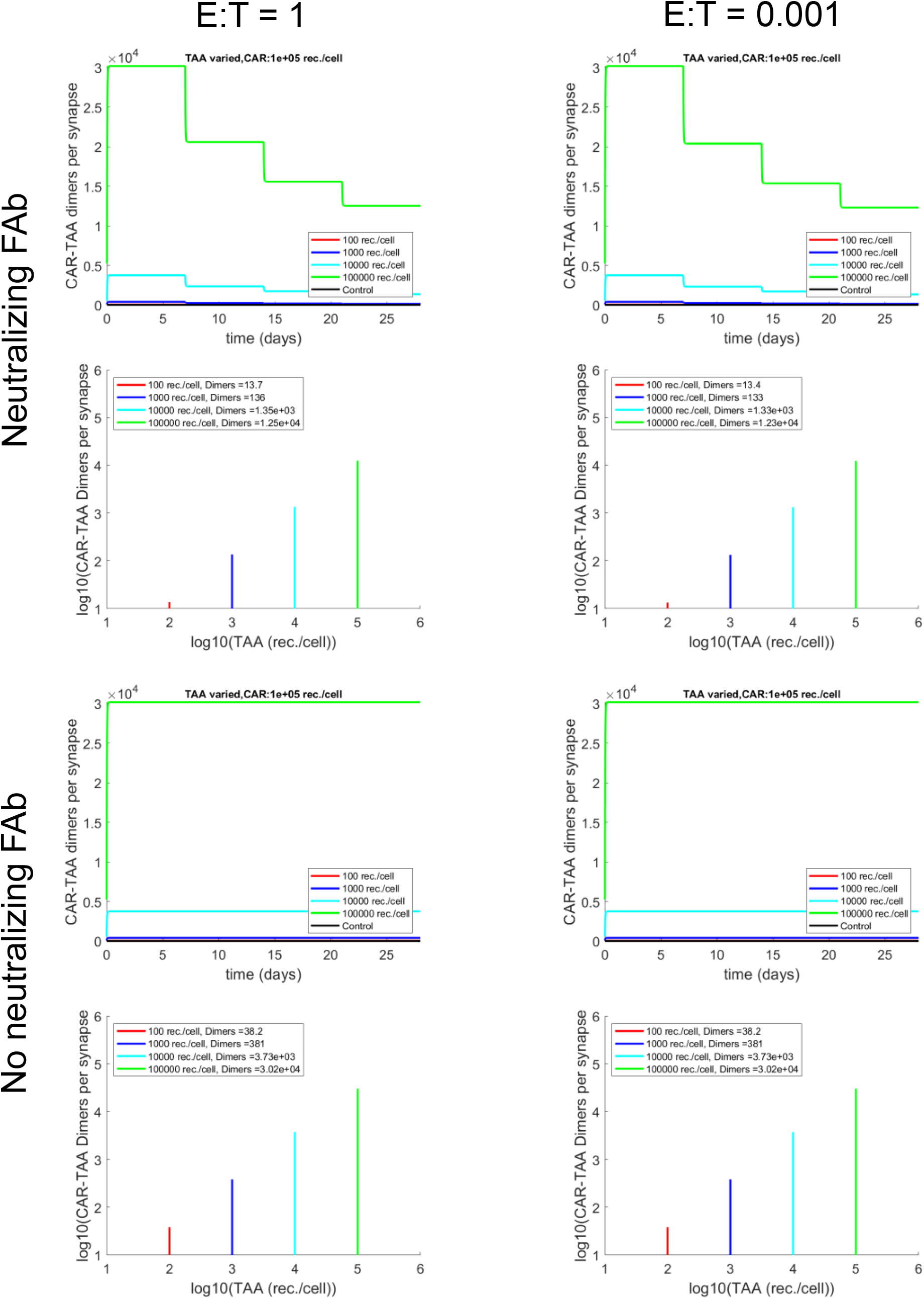
Inhibiting the CAR-T/TAA interaction with a neutralizing FAb that binds to TAA. Top panel: for different TAA expression levels, a FAb reservoir concentration (dose not deplete) of 2 nM was added starting on day 7, repeating every week, reaching 6 nM on day 21. Bottom panel: for different TAA expression levels, no Fab added. For each panel, Upper plots: time-dependent dimers per synapse for different TAA expression levels, Bottom plots: dimers per synapse at the end point in time for different TAA expression levels, Left column: E:T ratio of 1, and Right column: E:T ratio of 0.001. Here the avidity factor is Γ = 5.

## Supporting Information

### S1 Trimer formation rate for single synapse complexes assuming non-trimer species have the same concentration in and outside the synapse, i.e. extremely fast transport timescales

We’ve tested transport rates coupling the synapse and non-synapse regions which correspond to spread times of about 6s to over 2 hours. If we enforce extremely fast timescales of species redistribution, a given non-trimer species will have the same concentration inside and outside the synapse, when there are trimers (presence of bsAb) or none (no bsAb). Given this concentration constraint, we do not have to explicitly model transport in and out of the synapse, which will yield a slightly different form of the reaction rate for the trimer. To start, the total free TAA on the target cell of a synapse we denote as 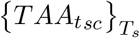. This implies that

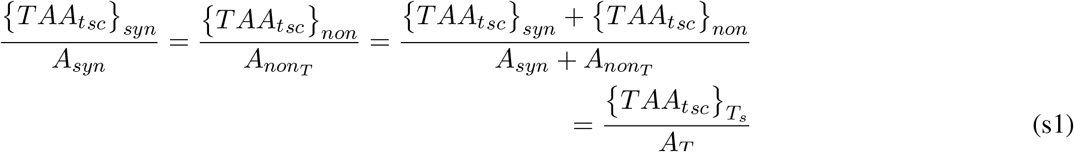

which implies 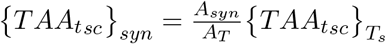. Likewise, the total IAA bound to bsAb on the immune cell of a synapse we denote as 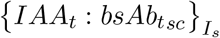. Our concentration constraint implies that

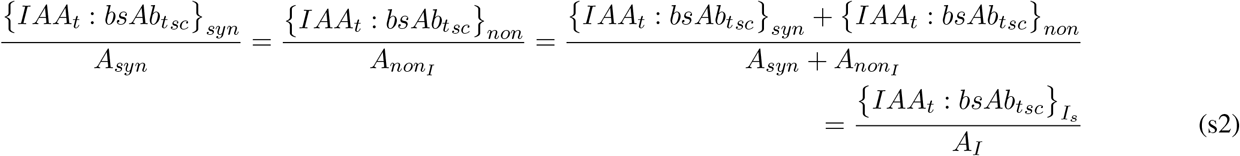

which implies 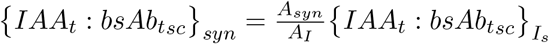. Given these assumptions and resulting relationships, the term for the reaction rate of a TAA binding to a bispecific antibody bound to an IAA (IAA-bsAb dimer) can be written as

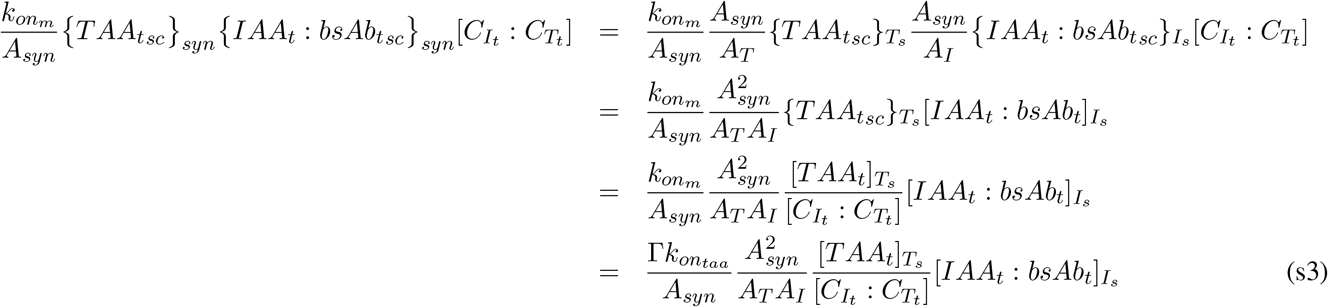

where we have represented trimer formation rate with total concentration of each species, i.e. the sum of the synapse and non-synapse concentrations. This alleviates the need to explicitly model transport terms as well as the synapse and non-synapse regions.

### S2 Application of the synapse model to Chimeric Antigen Receptor T cell therapies and general immune cell/target cell interactions

Because our synapse model is general, we can easily apply the approach to other immune cell therapies such as Chimeric Antigen Receptor T cells (CAR-T). Like ICE, CAR-T has found success in liquid tumors such as multiple myeloma [31] but solid tumors present a tougher challenge [32]. Figure S10 pictorially presents how our model can be adapted to CAR-T from the formation of the synapse to the binding of CAR and TAA forming dimers (CAR-TAA). Other than dimer formation for the CAR T case versus trimers for the bispecific case, the synapse formation and activation through cross-linking are similar between the two modalities. Importantly, the form of Eq. 3 is applicable to the CAR/TAA 2D diffusion reaction to form a dimer. In addition, the derived transport rate relationships above are also applicable to CAR-T synapses.

As a simple TGI demonstration, we apply the same basic killing function to our CAR-T TGI model as we did for the bsAb TGI model in Figure 6 (see the CAR-T transient synapse model in Section S3 for its mathematical form). Note that for these simulations we kept the E:T ratio constant. In Figure S11, we present simulated results of the model. In the left set of plots of Figure S11 we vary CAR expression with a nominal TAA expression of 10^4^ receptors per cell. For the right set of plots we vary TAA expression for a nominal CAR expression of 10^5^ receptors per cell. For both cases as one increases expression, the TGI gets stronger because the number of CAR-TAA dimers increases as well. As with the bispecific antibody case, for CAR-T, we can also apply the form of the thresholded trimer killing function (Eq.12) but applied to CAR-TAA dimers for different threshold values for some simple TGI examples at constant E:T ratios (Figure S12).

In addition, we can inhibit the CAR/TAA interaction by adding a FAb that binds to the TAA as a means of neutralizing the CAR-T therapy (Figure S13). While we focus on neutralizing the CAR-T therapy in this simplex example, this modeling approach also applies to endogenous immune cells as well where the FAb inhibition approach is directly applicable, for example, to PD1 (programmed cell death protein 1) on T cells interacting with PD-L1 (programmed death-ligand 1) expression on cancer cells [20].

### S3 Model variables and equations

The variables in the model describing different molecular and cellular species are represented as concentrations (nM for molecular species and cells/L for cellular species). For ease of notation and clarity of model equations, we will put square brackets around each variable, e.g. [*X*]. A description of the model variables are given in the table below.

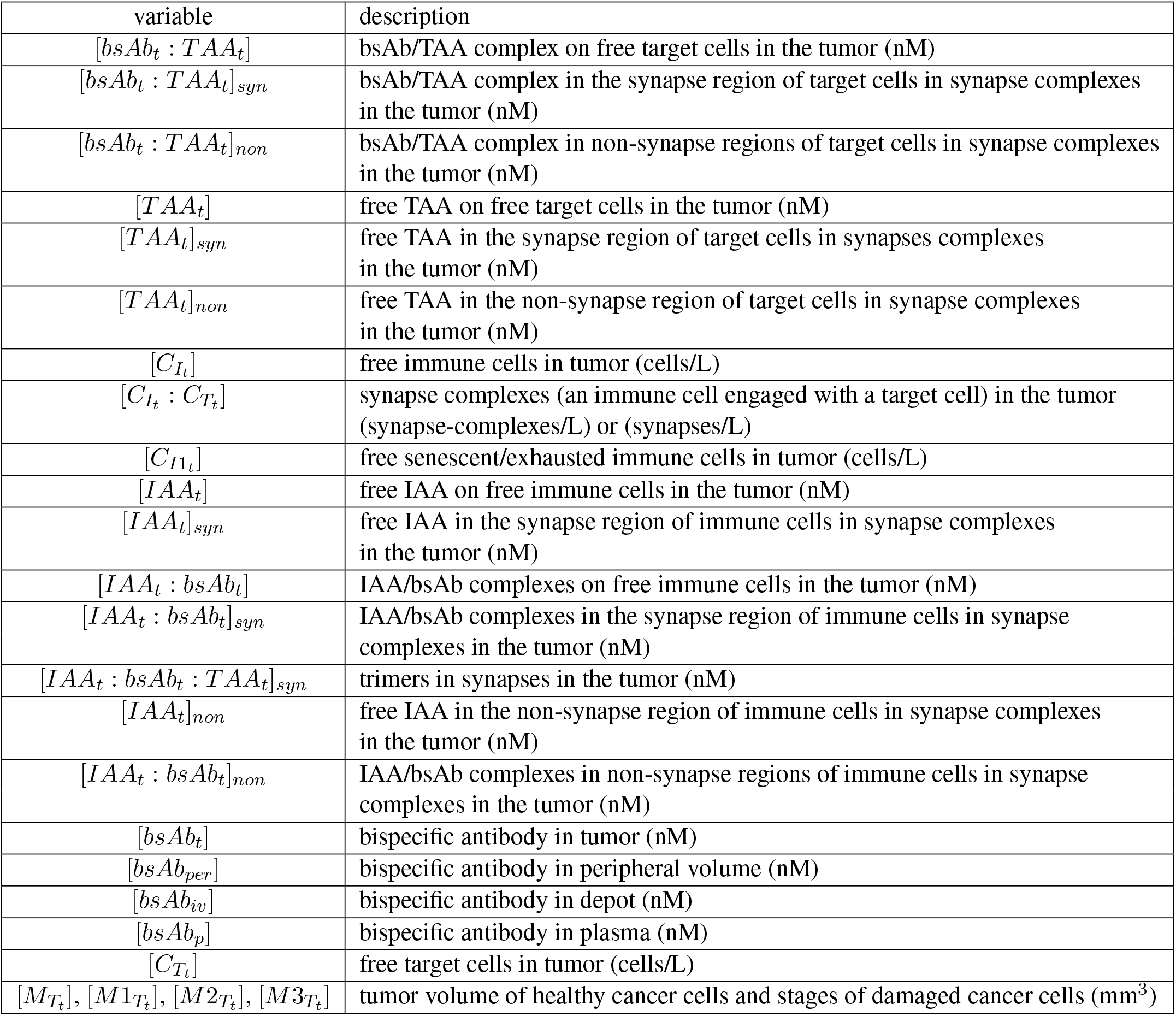

In the following sections we present 1) the equations for the model used in the initial studies of trimer formation (static populations, permanent synapses), 2) the equations for the model that incorporates a dynamic T cell population with transient synapses and a simple TGI model, 3) the equations used for the CAR-T examples.

#### S3.1 Synapse model where all synapses are permanent

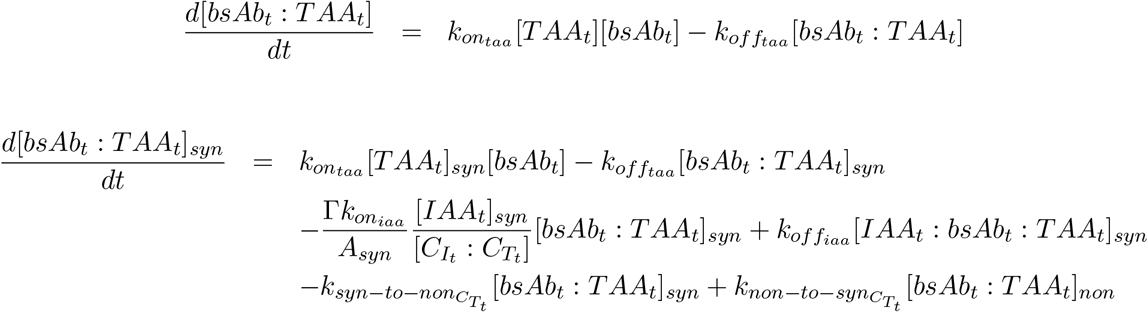

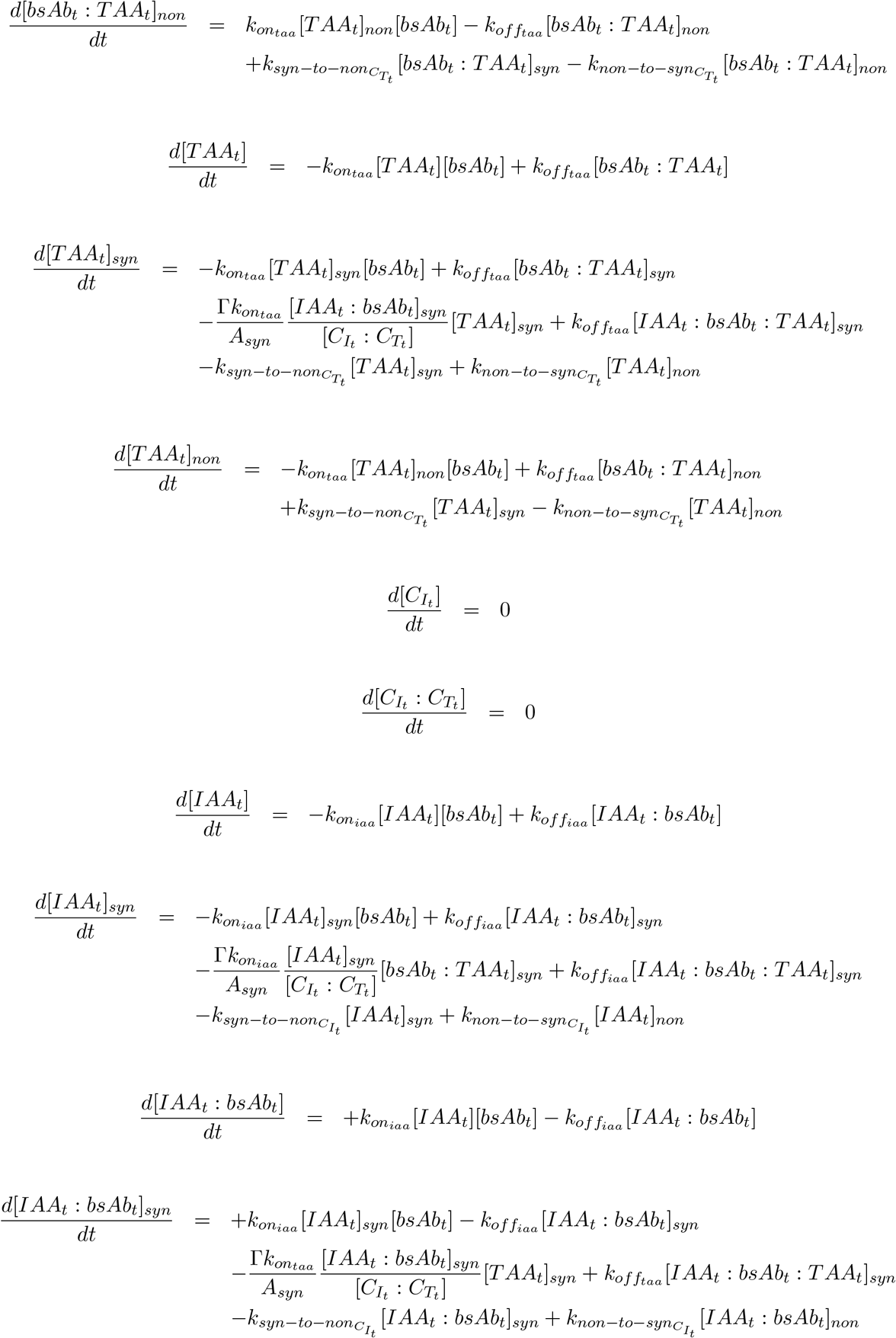

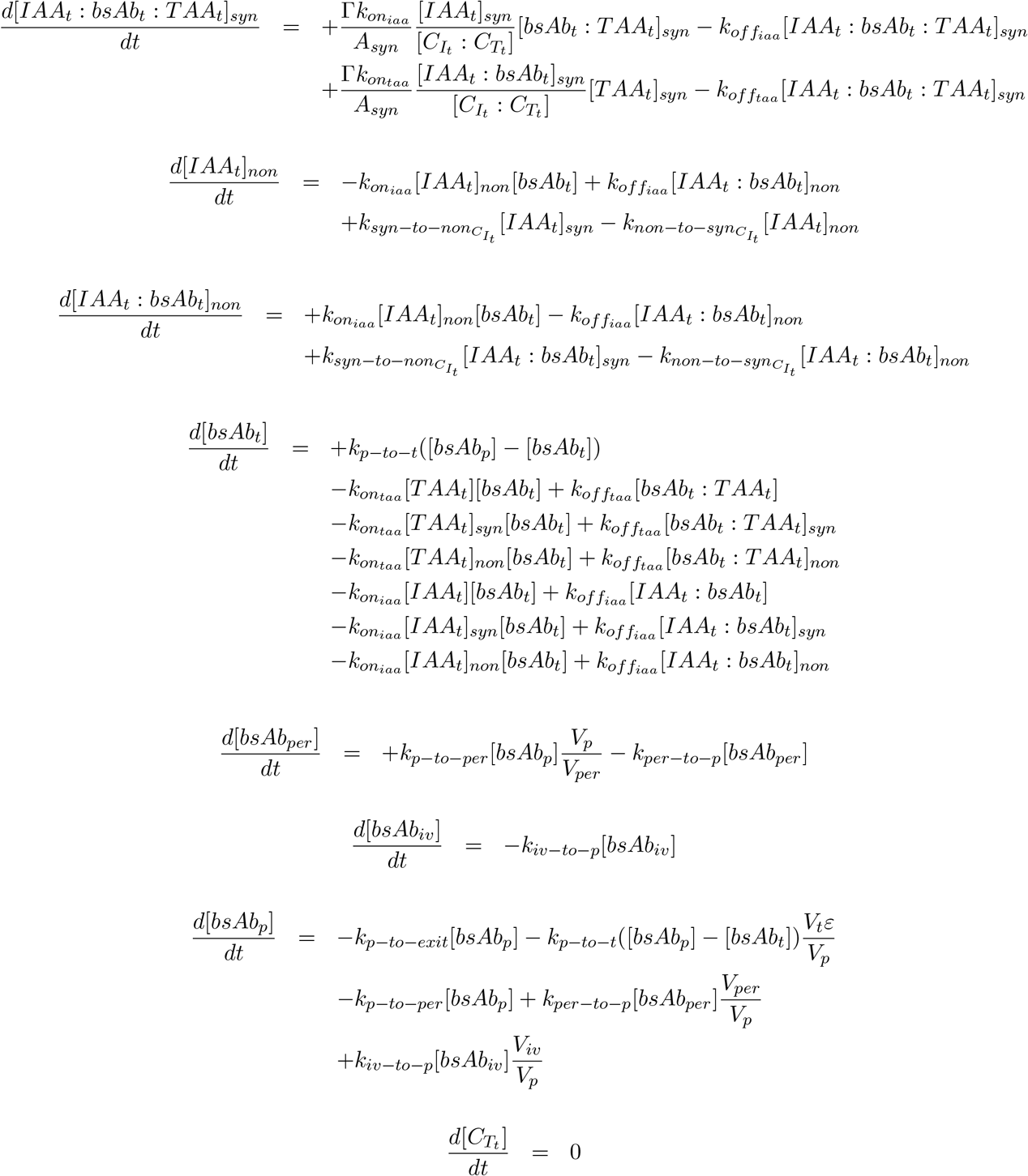

#### S3.2 Synapse model driven by a reservoir dynamic Immune cell population that form transient synapses

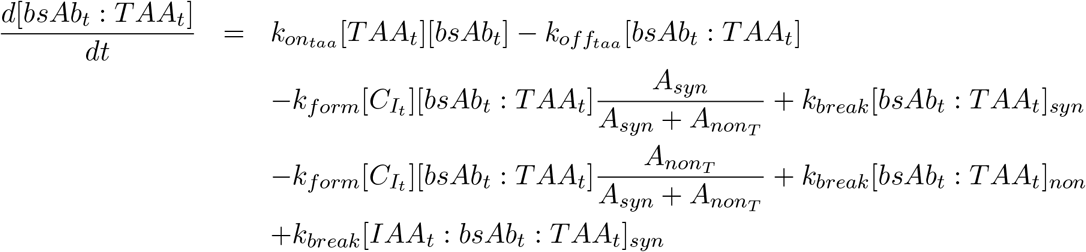

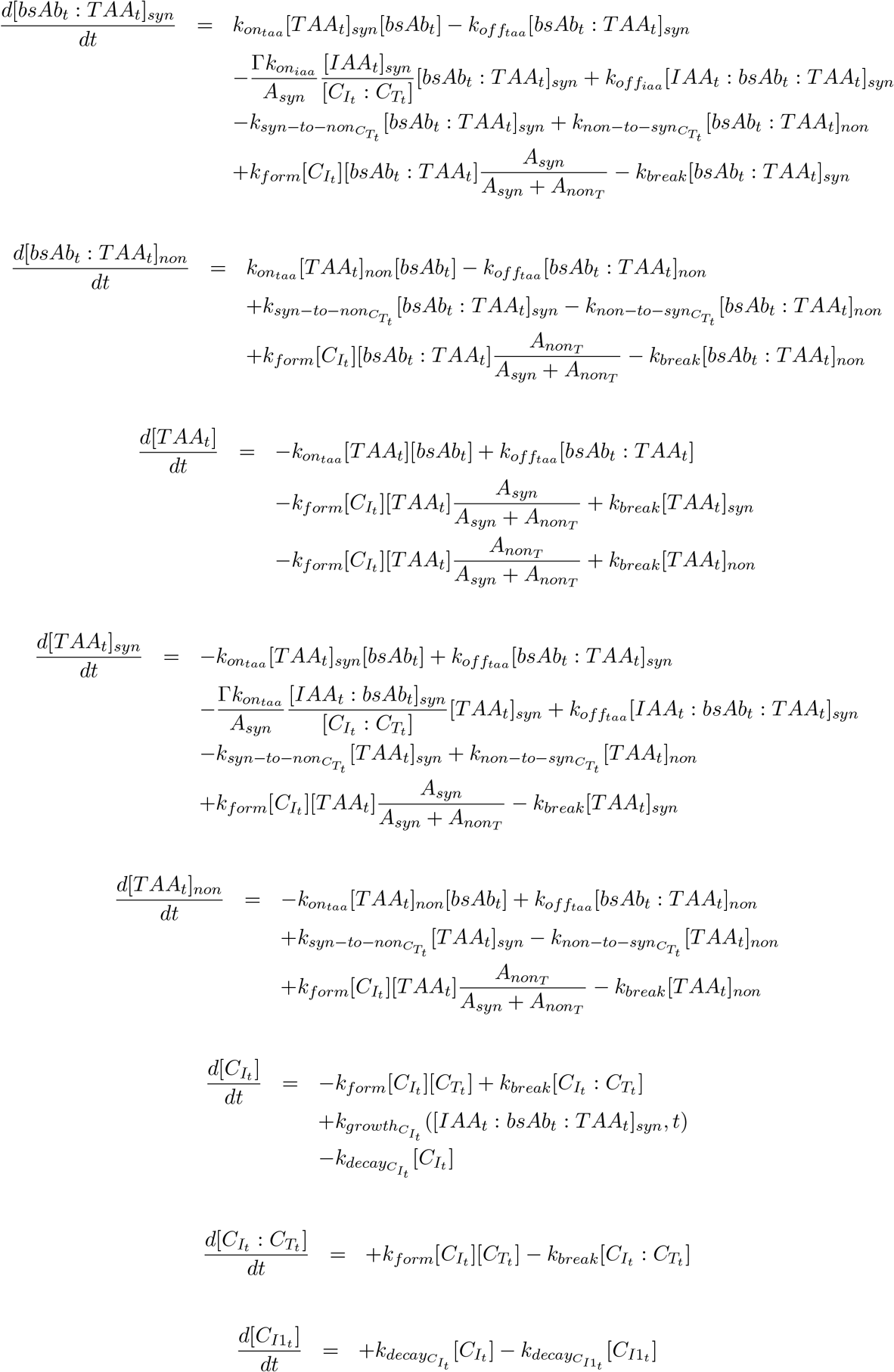

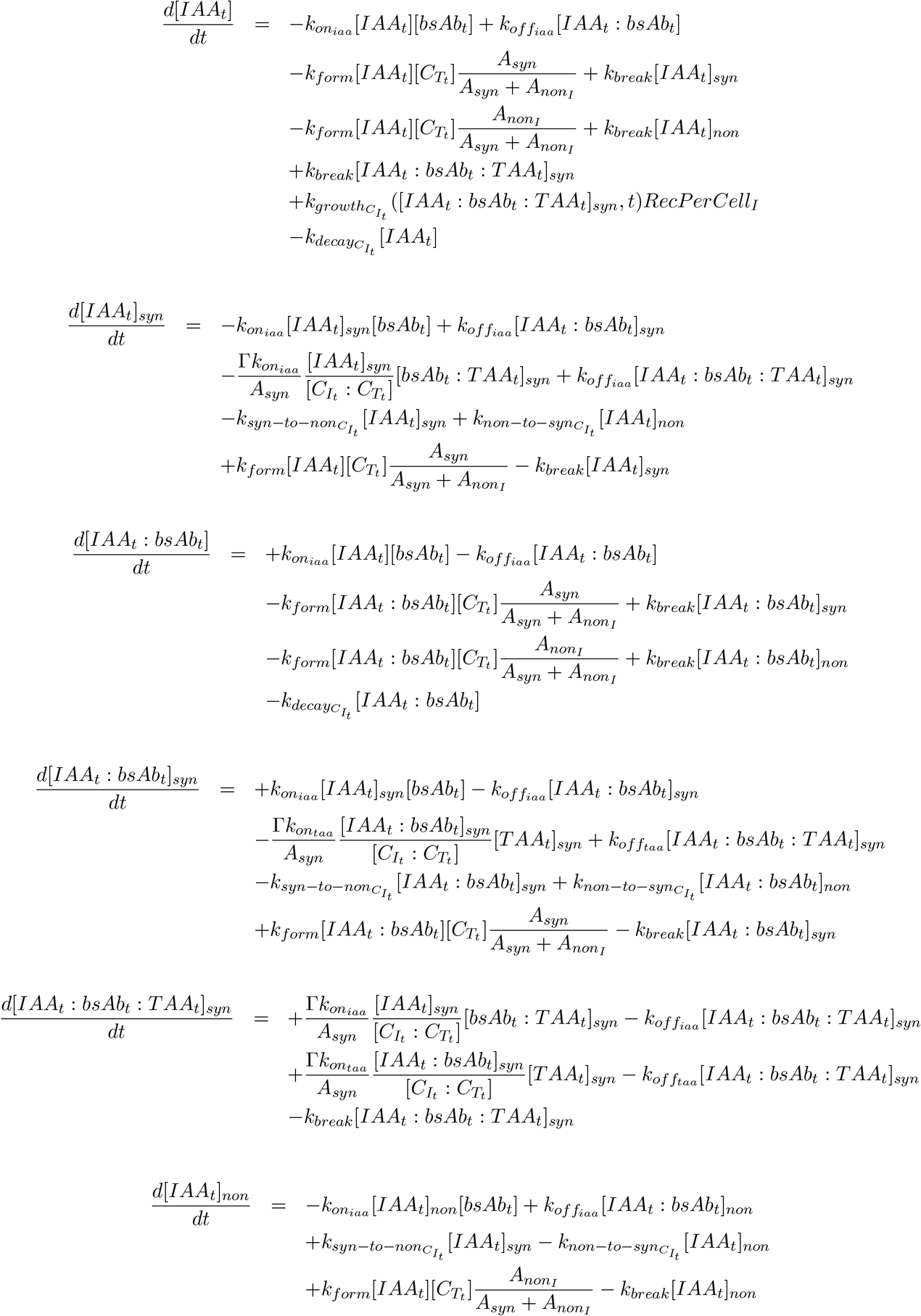

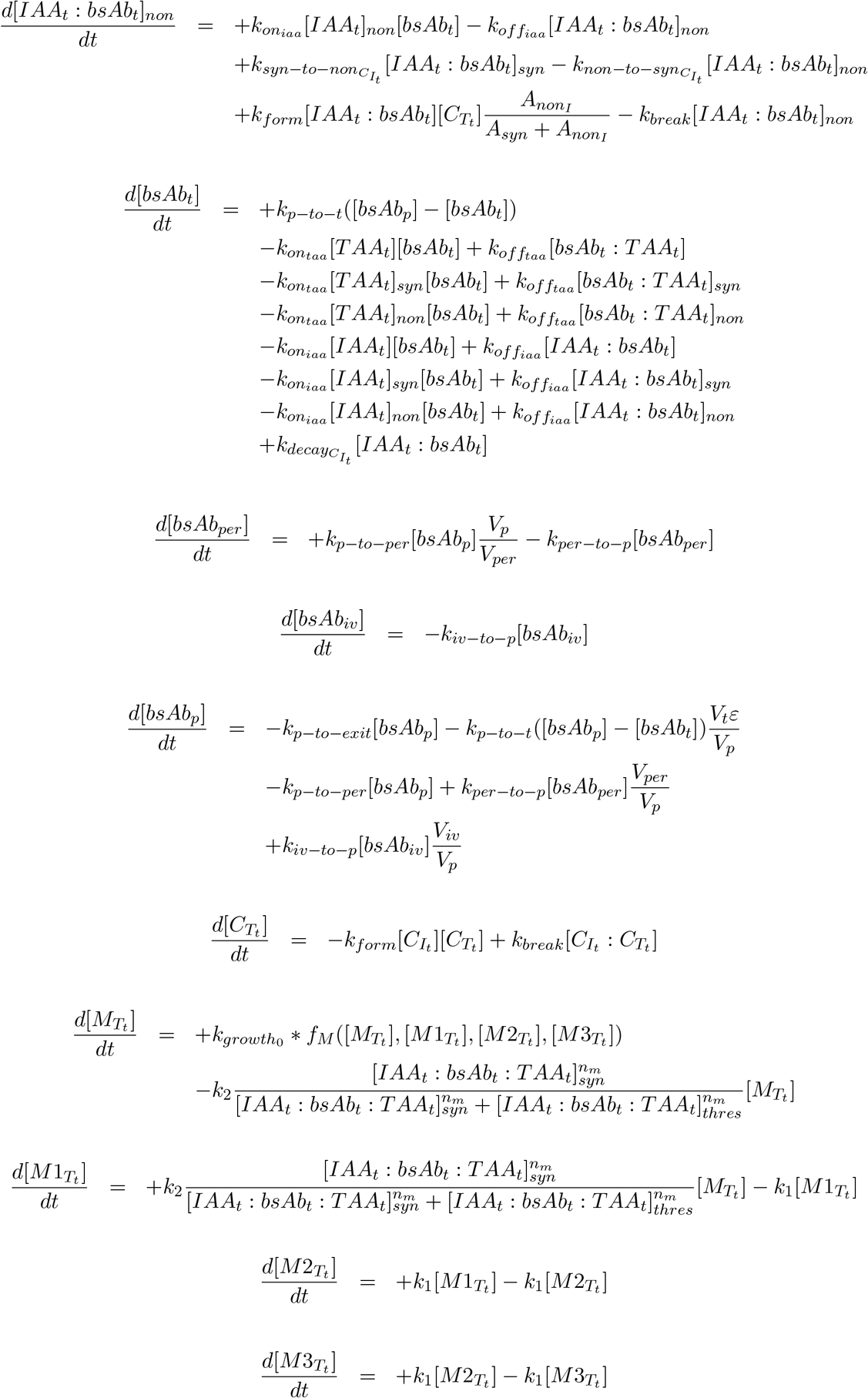

#### S3.3 CAR-T application of synapse model driven by a reservoir dynamic Immune cell population that form transient synapses

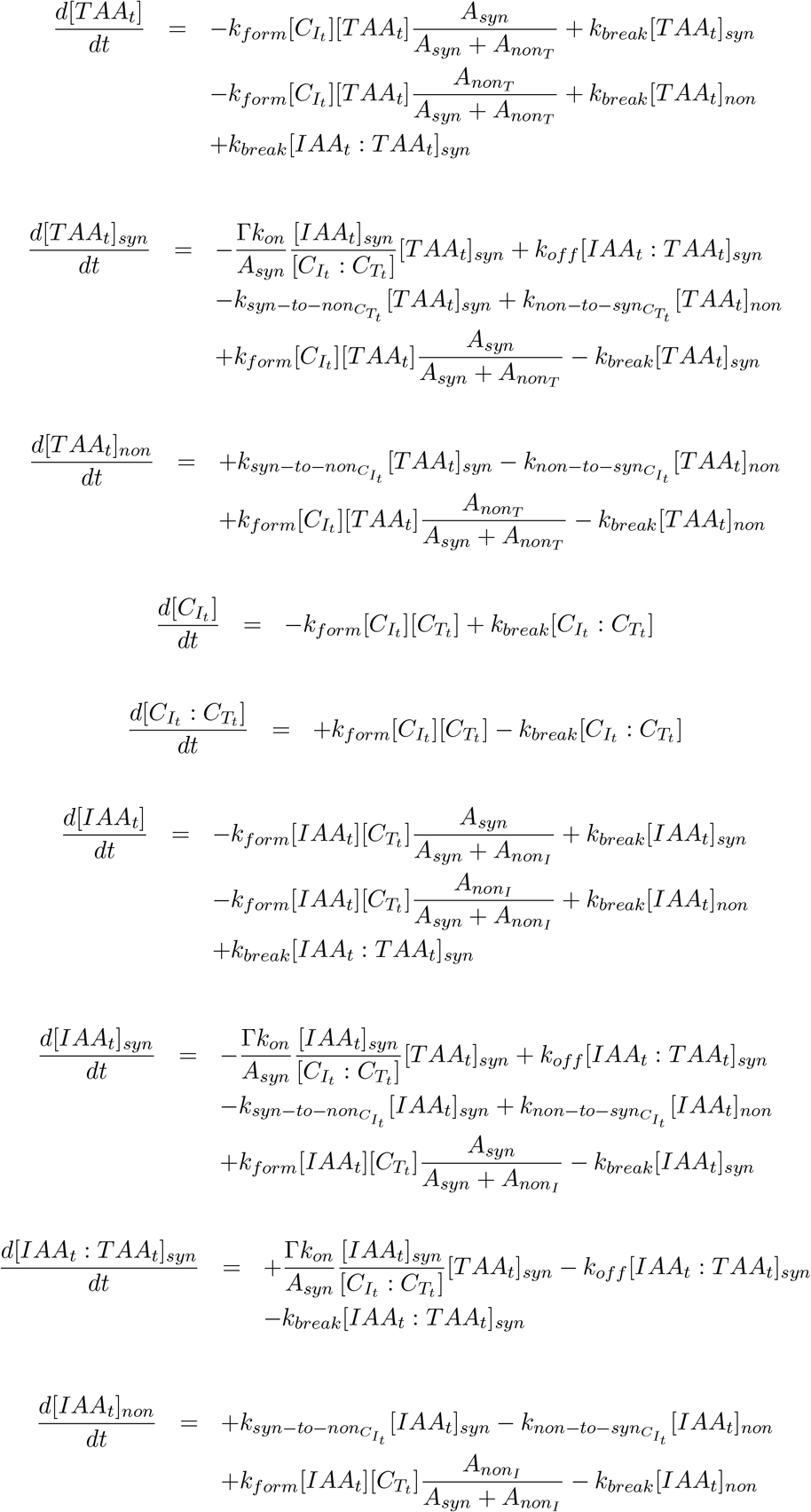

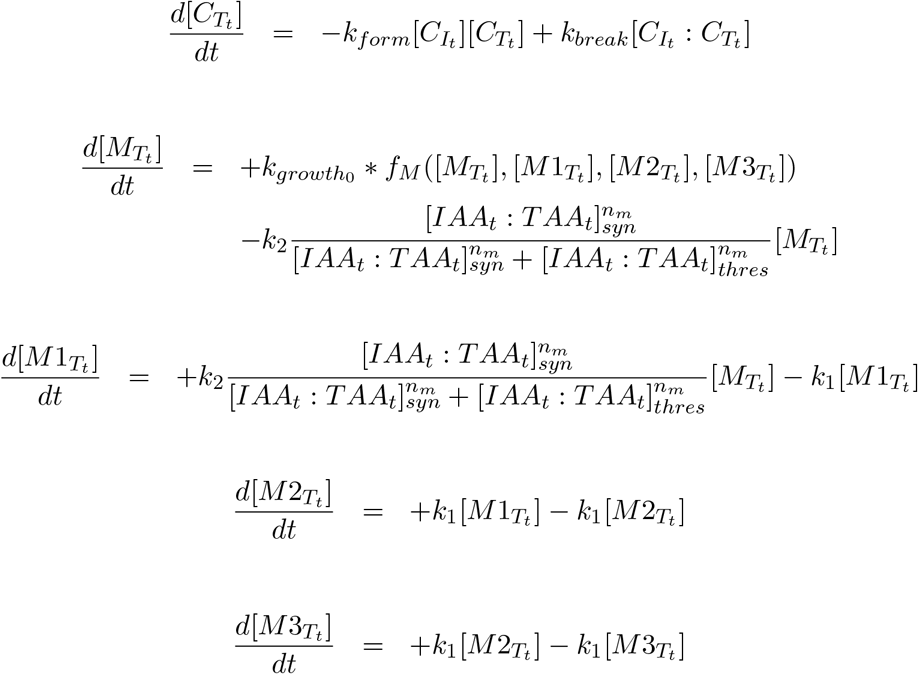

**Table S1:**
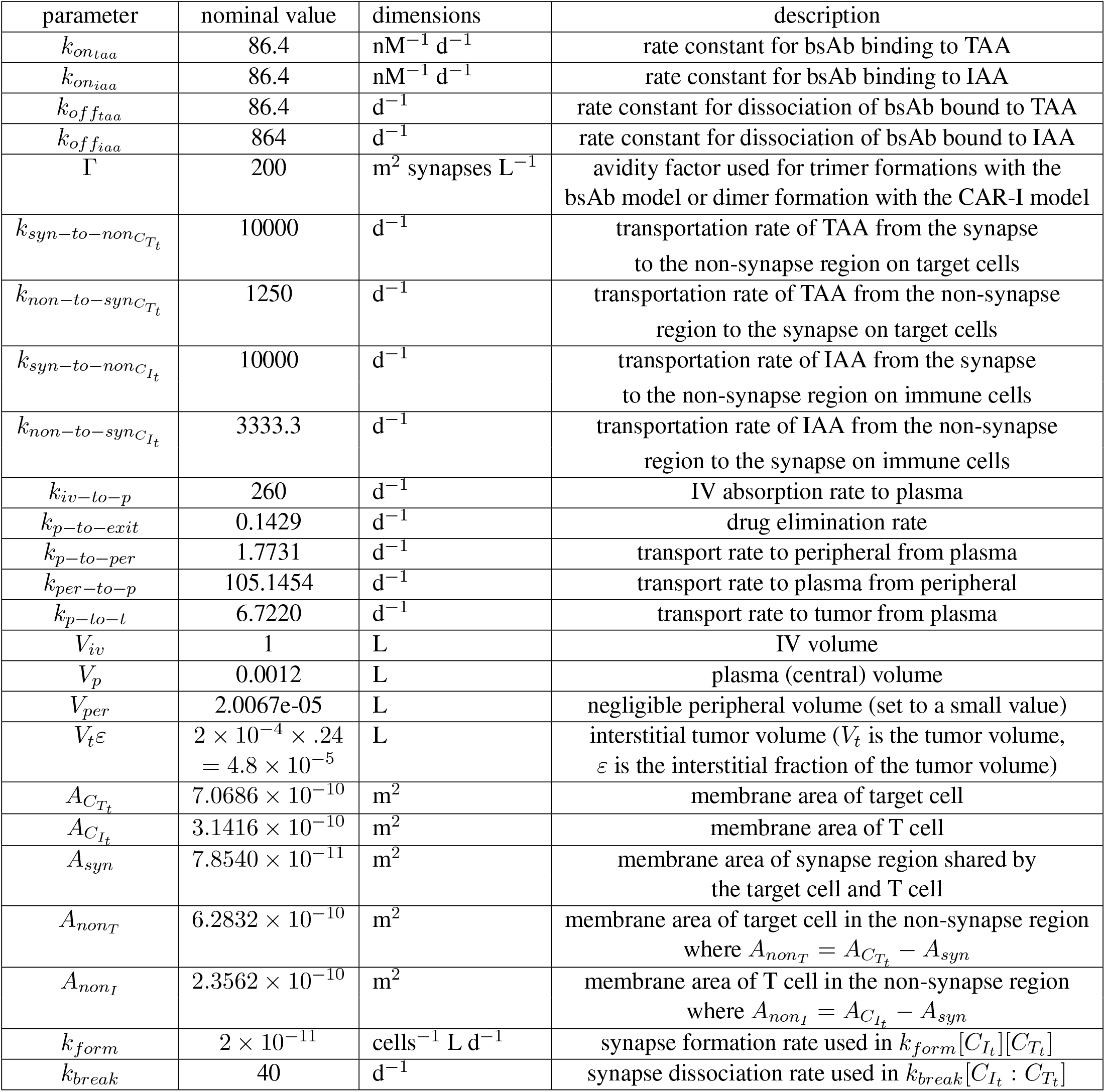
Simulation parameter values used in main Figures 1 through 5 and Supplemental Figures 1 through 7.

For all of the simulations, the total target cells and total T cells remain constant where individuals cells can be part of a synapse complex or not. For simulating trimer dynamics, the PKPD model used is effectively a 2 compartment model with the plasma compartment driving the tumor (site of action) and a negligible peripheral volume. For steady state trimer calculations, the model used is a one compartment tumor model (site of action) driven by a reservoir (constant) drug concentration. The tumor compartment has the all the same parameter values in both models. The total cells in the tumor volume is 2 *×* 10^7^ cells which equals the sum of target cells and immune cells in the tumor. The tumor interstitial volume is the same for all simulations. The total target cells equals the total tumor cells divided by (E:T ratio + 1). The total immune cells equals the total tumor cells divided by (1-1/(E:T ratio + 1)). This ensures the proper E:T ratio. Note that the tumor volume and its PK properties reflect that of a small vascularized xenograft tumor in preclinical mouse models.

